# Investigating local negative feedback of Rac activity by mathematical models and cell motility simulations

**DOI:** 10.1101/2025.05.05.651928

**Authors:** Jupiter Algorta, Jason P. Town, Orion D. Weiner, Leah Edelstein-Keshet

## Abstract

For polarization and directed migration, cells use a combination of local positive feedback and long-range inhibition. We have previously used mathematical models to show the ability of this core circuit to regulate directed cell movement. However, this wave pinning model lacks important additional feedback circuits, including the recently demonstrated local negative feedback from Town and Weiner. Here we extend our models to investigate the consequences of this additional link on cell physiology. We model responses of neutrophil-like HL-60 cells to spatially-controlled optogenetic stimulation of PI3K, leading (via PIP3) to Rac activity. We sequentially build up and investigate partial differential equation (PDE) models of the key Rac, Rac-Inhibitor, and PIP3-Rac-Inhibitor circuits. We fit model parameters to temporal and spatial (cell trajectory) data. Cell shapes, motility, and responses to stimuli are modeled in 2D cell-based simulations, with PDEs for Rac and the other regulatory components solved along the cell edge. We demonstrate that the ability of modeled cells to respond to temporal as well as spatial features of guidance cues depends on the addition of the local negative feedback circuit. Furthermore, the local Rac inhibitor improves the ability of modeled cells to respond to noisy or dynamic extracellular gradients. Our work demonstrates how local negative feedback enhances dynamic polarity and gradient sensing in migratory cells.

## Introduction

To navigate to sites of injury and infection, neutrophils have to polarize, align with shallow, noisy gradients of chemical signals, and undergo directed migration. It is well-known that intracellular components, including the GTPase Rac, play a crucial role in initiating and maintaining cell polarization, and transducing external gradients into regulation of the actin cytoskeleton and leading-edge protrusion. The spatial patterning of Rac depends on interacting networks of positive and negative feedback cascades that enable the generation of a dominant front that can be oriented by external guidance cues. *In vivo*, dynamic spatiotemporal gradients of guidance cues and complex 3D environments require the cell to continually adjust its direction of motion, balancing robust polarity with flexible response to its surroundings.

In recent work, we (Town & Weiner (2023)) used computer-controlled spatial optogenetic experiments to test responses of neutrophil-like HL-60 cells to stimuli that mimic reorientation signals in vivo. Cells were first polarized and then stimulated locally (on either left or right, front or rear edges). The light stimulus activates phosphoinositide 3-kinase (PI3K) signaling, known to produce the phosphoinositide PIP3 that, in turn, activates the GTPase Rac, leading to local F-actin assembly and front-edge protrusion. To determine the effect of stimulus history on cell response, cells were first exposed to localized PIP3 stimulus, followed by a global (whole-cell) PIP3 stimulus. This resulted in a zone of Rac inhibition at the previously-stimulated site, suggesting a local Rac inhibitor. During motility, cells reverse their direction of migration when the stimulus is switched from local to global, suggesting a guidance scheme that measures the local rate of change of the stimulus (see **Figure I**). This surprising result led Town & Weiner to propose a Rac-inhibitor, downstream of active Rac, that builds up alongside Rac zones, to dampen further Rac activation in those locales.

**Figure I:**
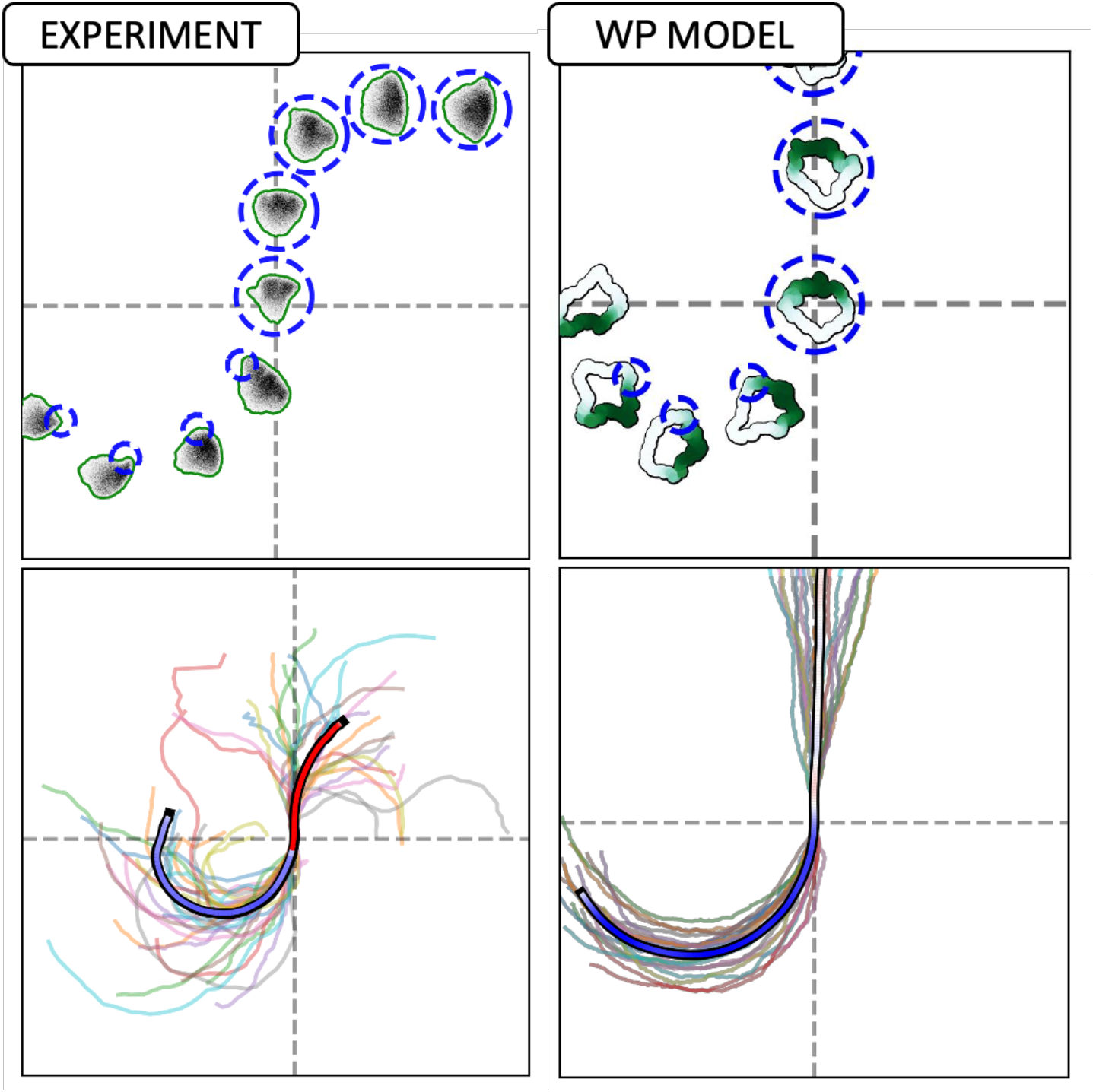
Exotic cell turning reversal is not captured by a basic model. Cells stimulated by local optogenetic inputs followed by global stimulation first turn in one direction, then reverse. Experimental data (left panels) show the directional behaviour and trajectories of cells, with snapshots (top left) and tracked positions (bottom left). The Wave-Pinning (Mori et. al., 2008, WP) model (right panels) reproduces the initial turning behaviour but not the reversal under global stimuli, showing cell shapes and trajectories (top right) and summarized paths (bottom right). In the lower panels, all individual cell tracks are shown (25 for the experiment, 20 for the model), with the average trajectory overlaid and colour-coded by rotation direction: blue for counterclockwise (CCW), red for clockwise (CW), and white for no rotation. Experimental data adapted from Town & Weiner (2023). Notably, while experimental cells reverse and begin turning CW after global stimulation, WP model cells continue along a straight trajectory. Video shown in Supplemental Material: **Movie 1**

**Figure II:**
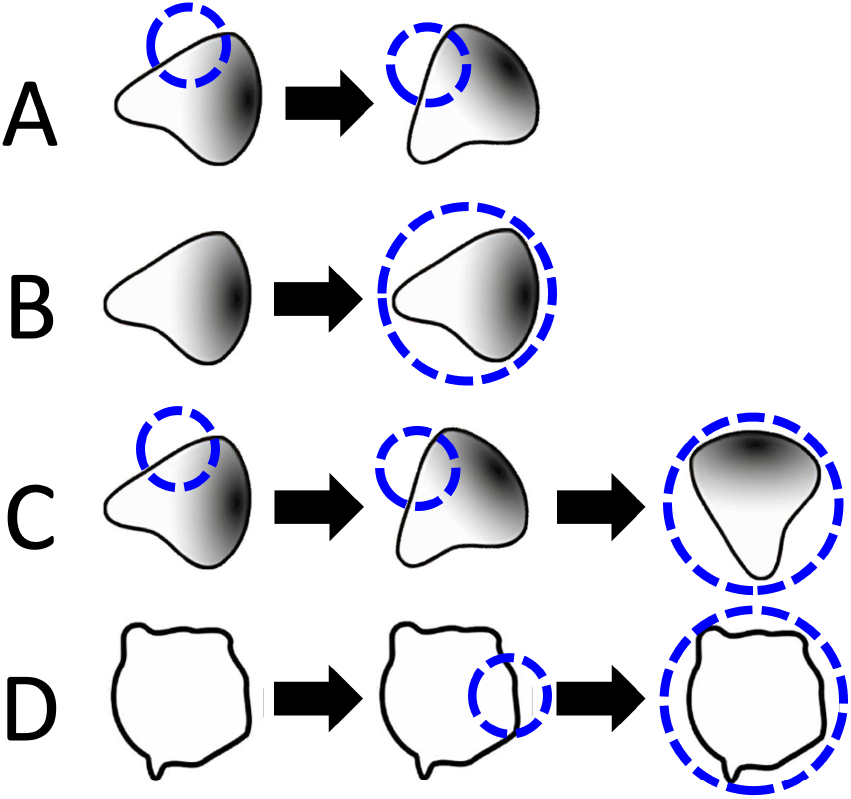
Stimulation protocols. The original stimuli in Town & Weiner included some combination of the above protocols. (A) Local stimulus on the side (or front or rear) of a polarized cell (B) Global stimulus of a polarized cell. (C) Local followed by a global stimulus of a polarized cell. (D) Treatment of a cell with latrunculin (to disassemble F-actin) followed by local and then global optogenetic stimulus.

The experiments in Town & Weiner (2023) provide compelling evidence for a Rac-promoted Rac inhibitor, while not yet identifying the relevant negative-feedback mediator. Yet, a few questions remain to be answered following those experiments:

1. Are there fundamental differences for how cells polarize and move with this circuit compared to our previously analyzed polarity circuit (Mori et al 2008)?
2. How does the proposed circuit that includes the Rac-inhibitor operate in situations where cells experience noisy gradients, or abrupt changes in time-dependent cues? Would the Rac-inhibitor allow for greater flexibility and gradient-following accuracy in the face of noisy, time-varying gradients?
3. What are molecular properties of the putative inhibitor? Can we infer its rate of diffusion (and hence its molecular weight), its rate of production downstream of Rac, and/or how effectively it inhibits Rac, based on the experimental data?

To address these questions and to rigorously test the proposed model in Town & Weiner (2023), we explore a sequence of mathematical models for Rac activity, in which we stepwise add its interactions with the putative inhibitor and with the upstream PIP3. Our models represent the time behaviour and spatial distribution of signaling components along the edge of a cell before and after optogenetic stimulation. The temporal and spatio-temporal model predictions are compared to the experimental data of Town & Weiner (2023), and in each case, parameter fitting is obtained by an optimization algorithm. Importantly, we also model the motion of the cell using a well-known method known as the cellular Potts model (CPM), where high Rac activity is linked to edge protrusion. In this way, we account for both intracellular Rac signaling and for the cell trajectories and responses to basic and “exotic” protocols.

### Experimental protocol

In Town & Weiner (2023), the authors express a combination of synthetic proteins in a neutrophil-like cell line. One pair of these proteins enable light-based control over a protein-protein interaction using the iLID system (Guntas, 2015). By localizing one light-sensitive protein to the plasma membrane and fusing its binding partner to biochemically active “cargo”, the authors were able to recruit the biochemical cargo to the plasma membrane using light, which can be spatially patterned to stimulate some or all of a cell. The biochemically active cargo in this instance was a protein domain known to bind PI3K (Inoue, 2008; Toettcher, 2011), which phosphorylates phosphoinositides and plays an important role in cell polarity. Another set of expressed proteins acted as biosensors, enabling real-time monitoring of stimulated biochemical activity and downstream consequences of that activity. By combining image processing and automated light patterning, Town & Weiner were able to create reproducible optogenetic stimulation protocols. Town & Weiner recorded cell trajectories, stimulation patterns, and biosensor responses, and these were used in this study to fit parameters to our models.

### Mathematical models and simulations

#### (I) Modelling Rac activity on the edge of a cell

To model cell polarization and reorientation, we initially used a simple reaction-diffusion model for active and inactive Rac shown in **Figure III, A**. This model, originally introduced by Mori et al. (2008) has the following main assumptions: (1) The total amount of Rac is taken to be constant (on the timescale of interest). (2) Inactive Rac diffuses faster than active Rac (serving as a global signal as it is depleted). (3) Active Rac promotes its own activation. In a suitable parameter regime, these ingredients suffice to promote and then stall a wave of Rac activity that resolves into a polar pattern, consistent with the front of a polarized cell (Mori et al, 2008). This partial differential equation (PDE) model is often called the “wave-pinning” (WP) model. The simplicity of the WP model, and extant mathematical analysis make it a convenient starting point for our investigation. In the sections that follow, we build on this foundation by introducing additional components to account for more complex behaviours

**Figure III:**
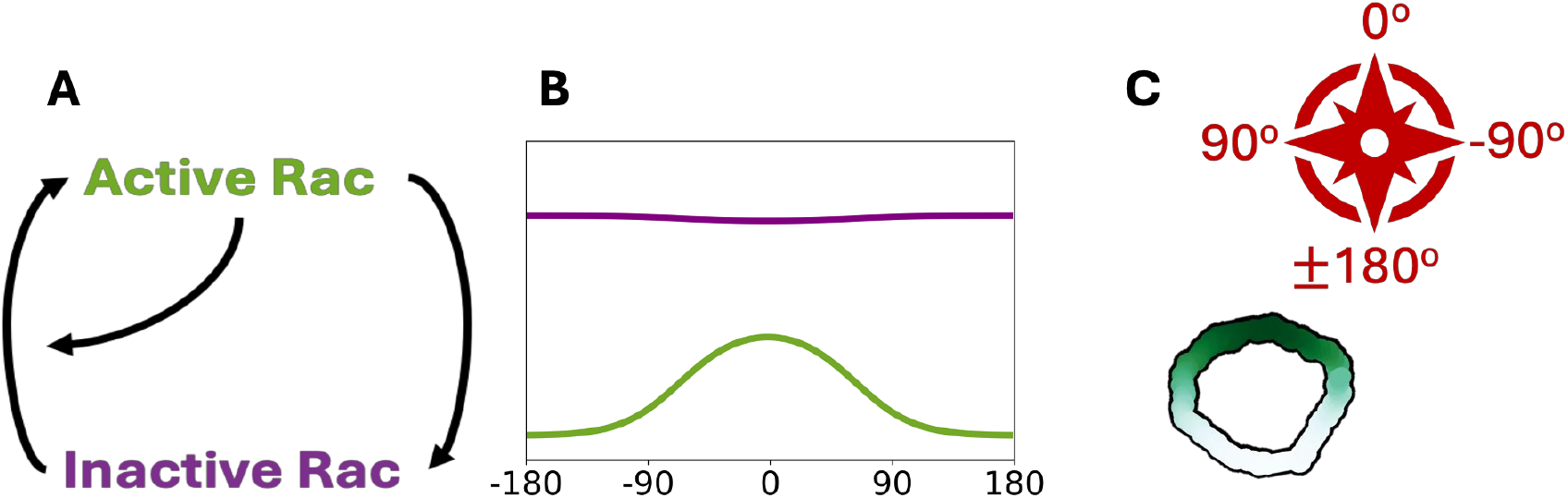
Geometry and basic model setup. (A) Basic set of interactions in the simplest model for Rac by Mori et al (2008) (“wave-pinning”) is formulated as a set of reaction-diffusion equations. (B) The model predicts the distribution of active (green) and inactive (purple) Rac along the cell edge, (a wraparound 1D domain, shown “unwrapped” in B). (C) A polarized cell is represented by the same nonuniform Rac activity distribution along its edge, with high Rac (dark green) defining the cell front. (Here the cell is polarized Northward, i.e., active Rac zone peaks at 0_o_.) The shape of the cell and its trajectory is simulated in the software Morpheus. The “compass” indicates the lab frame of reference, with 0_o_ corresponding to the Northwards direction. In simulations, the cell edge is shown as a wide band for easier visualization of the Rac distribution.

For a multiscale simulation of cell motility that encompasses intracellular signals, we used Morpheus (Starruß et al., 2014), an open-source simulation platform based on the cellular Potts model (CPM). Morpheus allows us to implement the distribution of Rac, its effector(s) and/or upstream signals along a 1D periodic representation of the cell edge. The distribution of active Rac is linked to edge protrusion and hence dictates the cell’s direction of motion (**Figure III, C**). The cell’s “front” is defined as the region with the highest concentration of active Rac. Further details on model equations and implementations are provided in the Supplementary Information.

In contrast to other models for cell polarization (Meinhardt, 1999; Otsuji et al., 2007; Neilson et al 2011a, 2011b), the WP model does not follow the typical Turing activator-inhibitor template, nor the local-excitation-global-inhibition (LEGI) structure (Levchenko & Iglesias, 2002). Rather, it is more closely associated with a “substrate depletion” mechanism, where the reservoir of inactive Rac is used up during Rac activation. Furthermore, WP operates outside the usual “Turing pattern forming” regime, meaning that WP permits both resting (uniform) and polarized cell state to coexist, so that cells can respond to signals that are sufficiently large while avoiding a response to small noisy inputs.

The WP model has several parameters that determine its behaviour: the basal (*k*_*0*_) and the autocatalytic rates of Rac activation (γ), the rate of inactivation (d), and rates of diffusion of the active and inactive forms (*D*_*u*_, *D*_*v*_ respectively). Rac autocatalysis is represented as a saturating (Hill) function of the Rac activity, with a half-max parameter (*K*): the Rac level *R = K* leads to enhanced Rac activation rate of γ/2. The values of these parameters were initially set as in Mori et al (2008) to investigate qualitative model predictions, and then fit to our experimental data, as described below.

#### (II) Modelling the optogenetic stimulus

To model the effects of external stimuli, we incorporated an additional rate of Rac activation at the site on the cell edge affected by the stimulus. **Figure IV** illustrates two types of stimuli used in this study:

**Figure IV:**
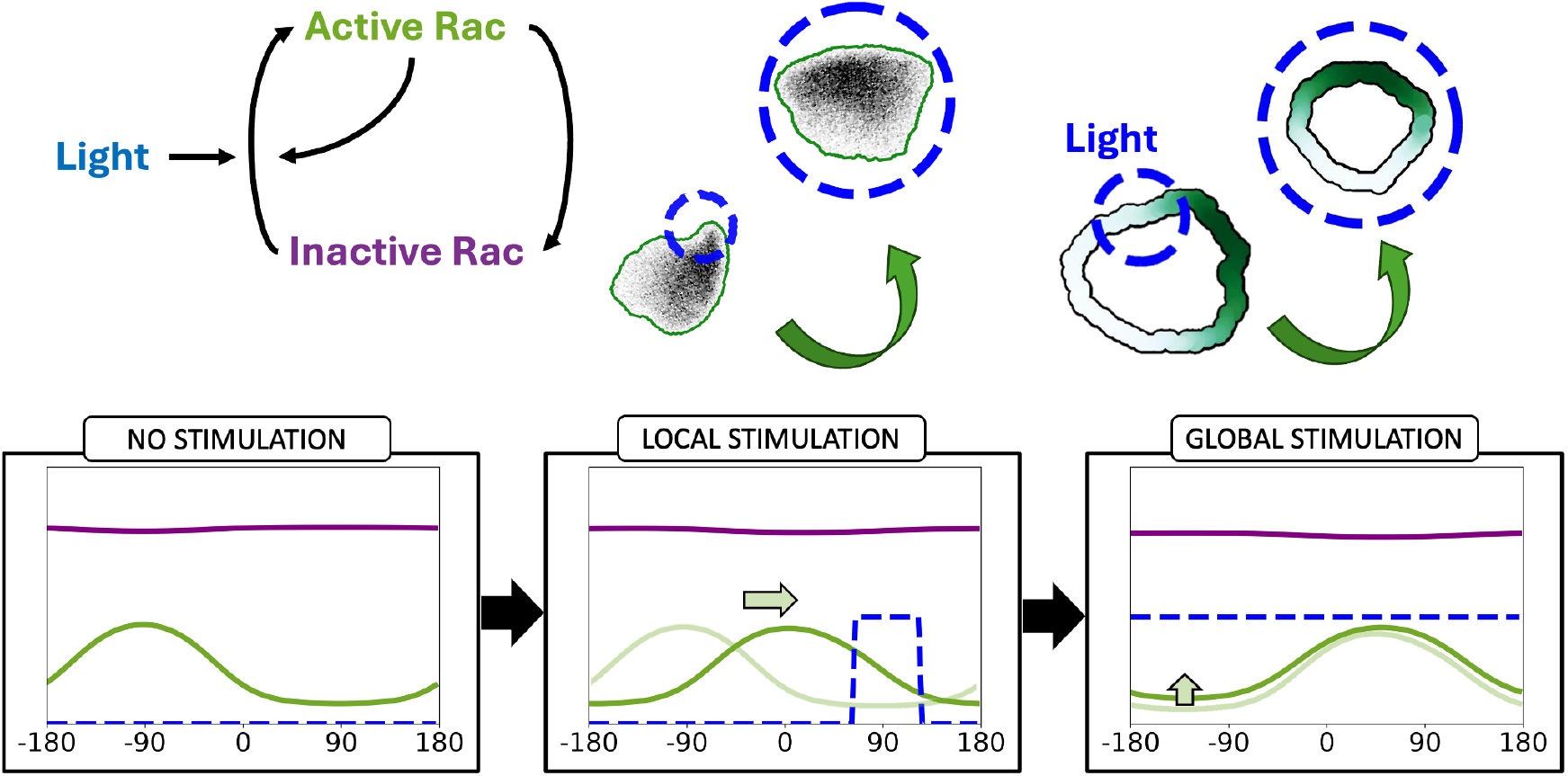
Modelling the optogenetic stimulus. The light stimulus is assumed to increase the rate of Rac activation in the models (see SI/Appendix). The location of the stimulus is represented by the dashed circle (top). We track the location of the stimulus and the model components in the 1D profile (bottom row) as well as the moving cell edge simulation. Blue: light stimulus, green: active Rac concentration, purple: inactive Rac concentration along the cell perimeter.

1. Local stimulus that activates Rac in a specific region of the cell edge.
2. Global stimulus that affects Rac activation across the entire cell edge.

The profiles of active and inactive Rac are shown (green, purple) in the bottom panels of **Figure IV**, under three conditions: no stimulus, localized stimulus, and global stimulus. The light stimulus is shown in blue.

## Results

### Result 1A: An elementary polarity model (WP) can account for simple behaviours

We first tested the simple WP model of Mori et al (2008) and its predictions for the polarity and motion of a cell exposed to basic local optogenetic protocols shown in **Figure V**. Recall that in experiments (labelled “Experiments” in **Figure V**) the local stimulus at the rear of a polarized cell will cause the cell to reorient and make a U-turn, while two local stimuli at opposite sides of the cell will result in one or another direction (right or left) “winning”, and cause the cell to turn right or left. We found that the WP model easily accounts for such behaviour.

**Figure V:**
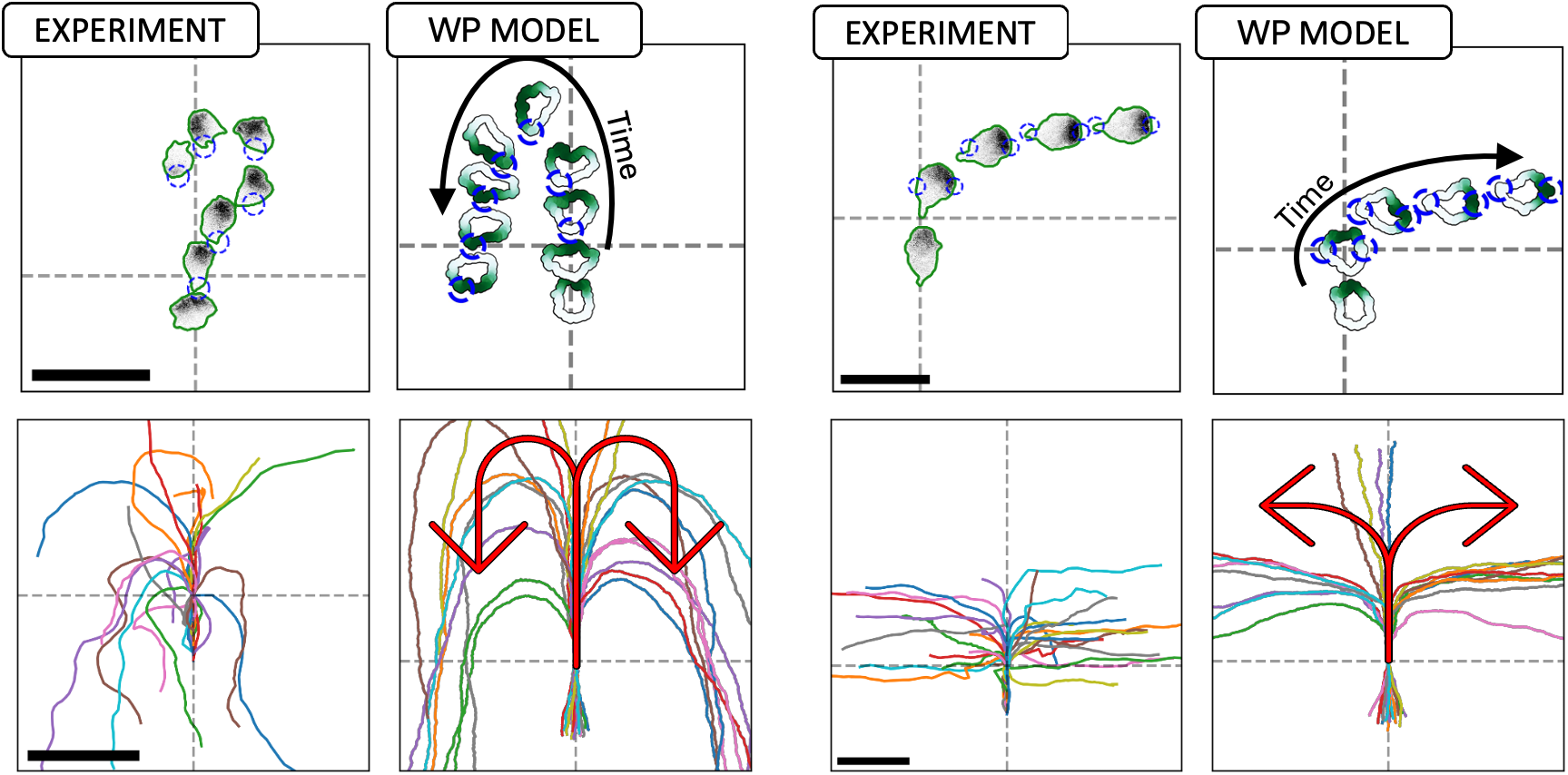
The WP model predicts cell responses to simple stimuli. The simplest model for Rac (WP) can satisfactorily account for cell motility responses to basic local optogenetic stimuli. Left two columns: experimental data and model predictions for local rear stimulation; the cells make U-turns and move in the reverse direction. Right two columns: competing two-sided local stimuli; The cell selects one or the other side and turns. Scale bar: 50μm. This demonstrates that, qualitatively, the WP model can reproduce the observed experimental cell behaviour for basic stimuli. The dashed axis indicates where the stimulation started. Video shown in Supplemental Material: **Movie 2**

**Figure VI:**
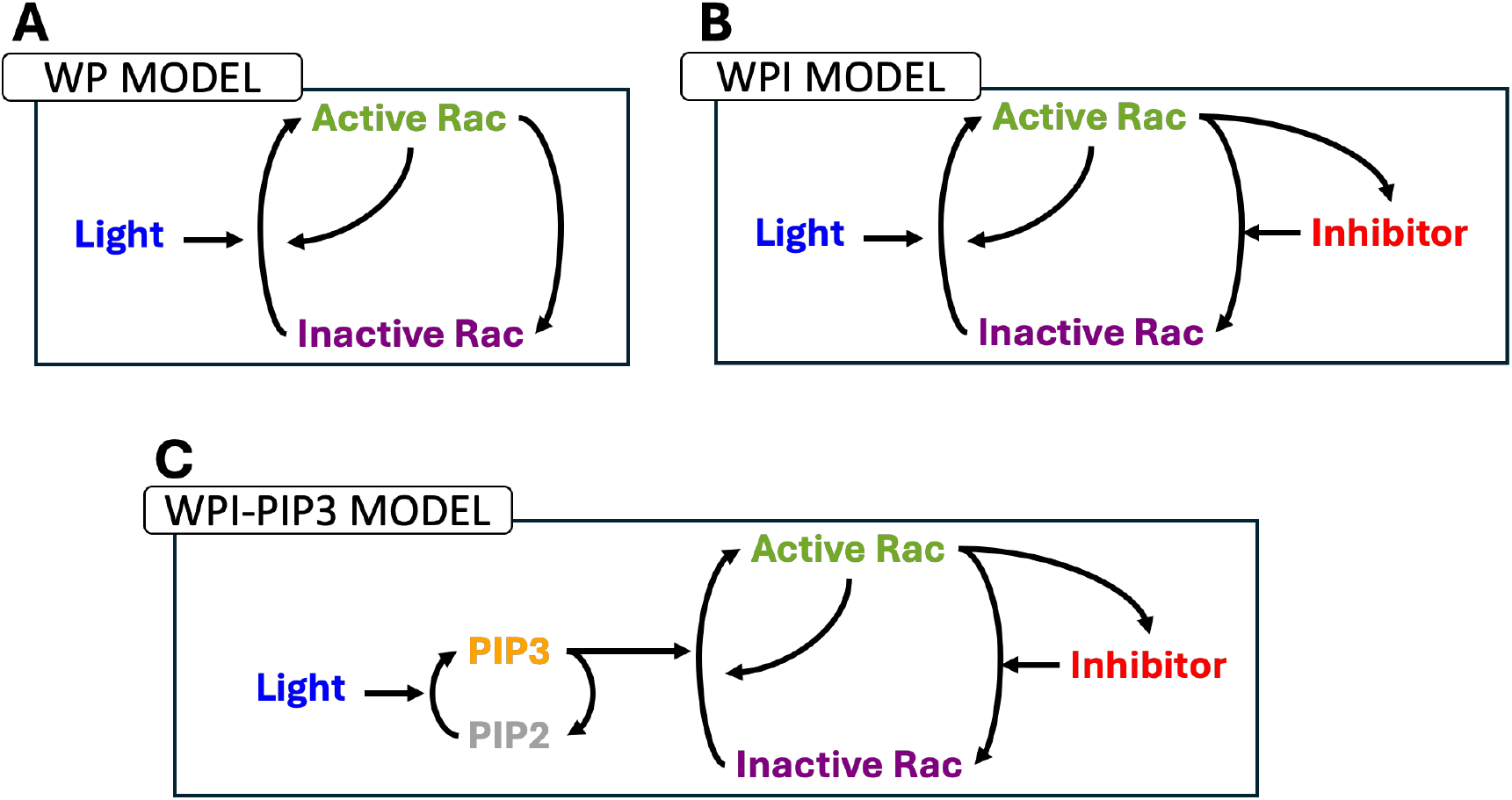
Three competing models. (A) The simplest model for Rac activation follows the “wave-pinning” (WP) polarity model of Mori et al (2008). (B) A Rac-associated inhibitor, proposed by Town & Weiner (2023) is included in the WPI (Wave-Pinning with Inhibitor) model. (C) The role of PIP3 was also considered in the WPI-PIP3 model. All models can account for simple response to single stimuli, but only the WPI-PIP3 model can also account for the behaviour observed in the reversal experiment.

### Result 1B: The behaviour predicted by the basic (WP) polarity model does not replicate the “local-then-global” stimuli experiment

We simulated the basic WP model with the protocol in which a local optogenetic pulse on one side of the cell is followed by a global stimulation of the entire cell. As shown in **Figure I**, the WP simple model fails to account for the observed reorientation of the cell (reversal of direction of turning). Town & Weiner (2023) proposed that this behaviour was due to the action of a local Rac-inhibitor downstream of Rac.

To test the idea of a Rac inhibitor, we revised the model, as shown in **Figure VI, B**. We added a PDE for the inhibitor (*H*), assuming that it is produced at a rate proportional to the level of active Rac, and that it decays with first-order kinetics. We also assumed that the inhibitor diffuses with some rate *D*_*H*_. Before examining the predicted cell trajectories, we asked what parameter values for the model(s) would be consistent with time-dependent data in Town & Weiner (2023).

### Results 2A: Model fits to the experimental time dynamics in latrunculin-treated cells

Before investigating the full responses of motile cells, Town & Weiner simplified the experimental system to showcase the temporal changes that take place in response to stimuli. To do so, they treated cells with latrunculin (Lat) to immobilize them and abrogate the effect of filamentous actin. Hence, in these experiments, the cell does not deform, and membrane tension plays no role in the dynamics of Rac. The time-dependent levels of PIP3 and active Rac were quantified experimentally by Town & Weiner (2023). Their data provide an opportunity to fit the model’s temporal behaviour before trying to understand the full spatio-temporal dynamics.

To fit parameters to the time-dependent models, we dropped the spatial (diffusion) terms, retaining ordinary differential equations for Rac (active and inactive forms), and its inhibitor. We used a parameter fitting routine (Price et al., 2006) to find best-fit parameters for each of the two models (WP, n = 58 and WPI, n= 87). To match the presentation of the experimental data, the model variables were scaled so that the unstimulated uniform active Rac level was set to “1.0” (baseline level) and its stimulated level treated as a “fold multiple” of that basal level.

We used a second improvement for fitting the data: the PIP3 data (rather than the optogenetic on-off timing) was used as a direct input to the models’ Rac activation rate. We also considered the heterogeneity of cells displayed in the data. Details of the fitting method are given in the SI.

### Results 2B: On-Off stimulus

Cells were exposed to a light stimulus that was turned “ON” instantaneously. The data displays features of adaptation (Chang & Levchenko, 2013), including a rise in Rac activity, followed by decay back to its basal level. Experimental observations (left), as well as model fits for WP (center) and WPI (right) are shown in **Figure VII**. We observe that the WPI is a better fit to the time dynamics than the WP model. In particular, the WPI model can account for the overshoot of Rac, whereas WP cannot.

**Figure VII:**
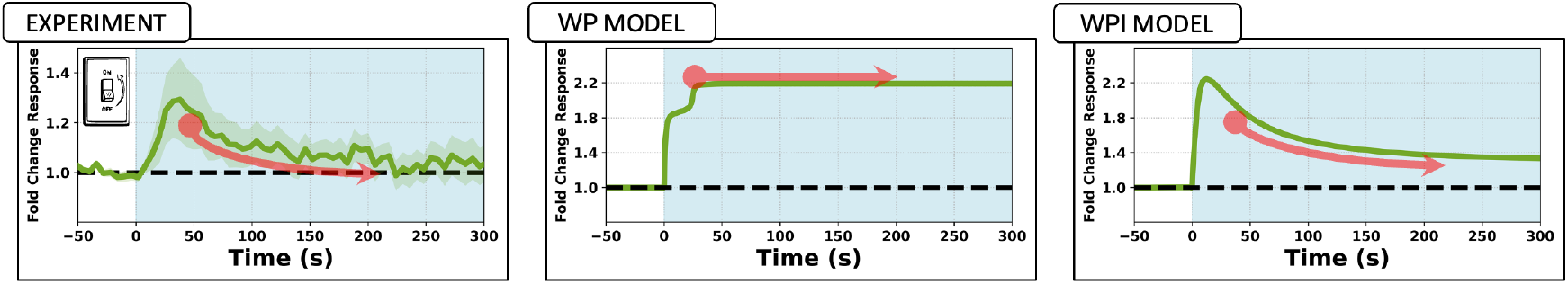
Experiment and model responses to stimulus turned ON. Active Rac fold change responses in green. Left: experimental data for an “ON” switch pulse stimulation of latrunculin-treated cells. Center: WP model predictions. Right: WPI model predictions. WPI is better at fitting the data and accounting for the decay following peak Rac activity. PIP3 levels quantified in the experiments were used as a direct input to Rac (as surrogate for the light stimulus) for the WP and WPI model.

### Results 2C: Double-pulse stimuli

A second type of stimulus used on Lat-treated cells was a double-pulse, consisting of two ON-OFF step functions, either Δt_1_ =120s or Δt_2_ =60s apart (top and bottom rows, respectively in **Figure VIII**). We compare the experimental data (left panels) in **Figure VIII** to model fits. It is typically observed that the second peak of Rac activation is smaller than the first peak, and significantly smaller if the pulses occur in rapid succession.

**Figure VIII:**
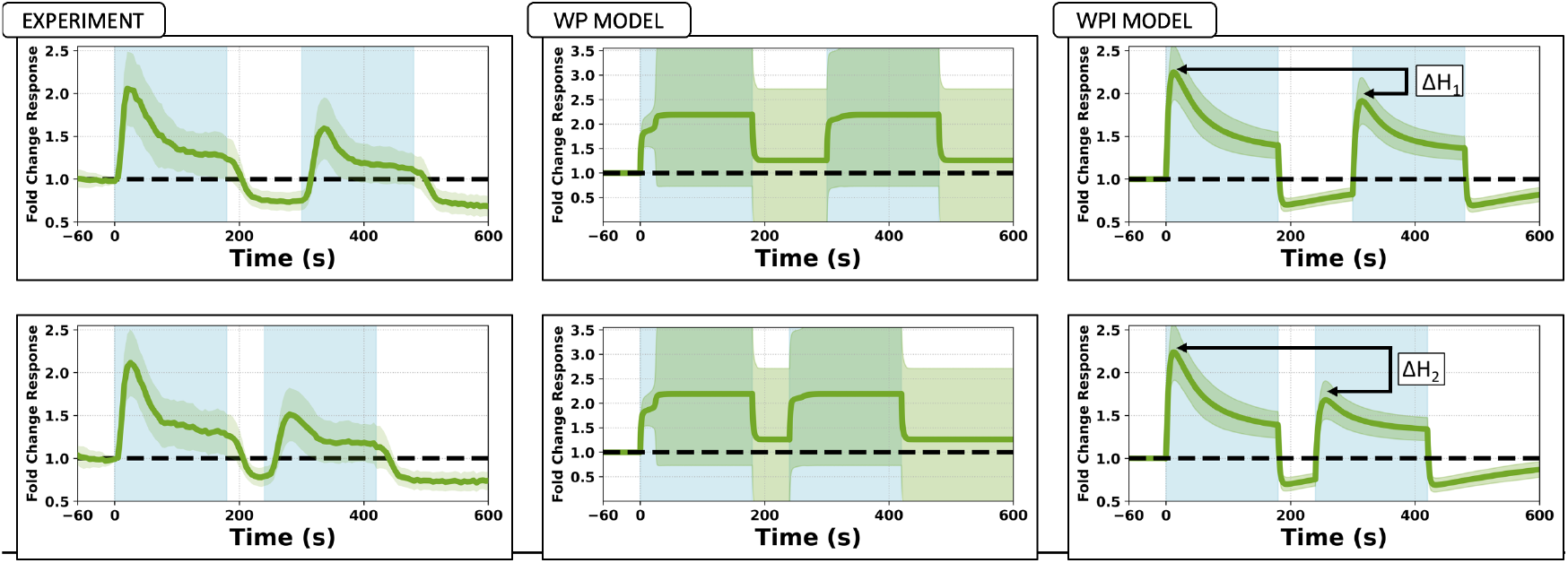
Double-pulse experiment. Data (Town & Weiner, 2023) (left) and model fits by the WP model (center, n=58) and the WPI model (right, n=81). Mean Active Rac (solid green curves) ±1 SD (green shading). Light blue represents stimulus ON phase. **Top:** a longer delay (Δt_1_=120s). Both models were independently fit to the data from the longer delay (Δt_1_=120s) experiment. **Bottom:** A short delay, (Δt_2_=60s) between stimuli. The results for the shorter delay (Δt_2_=60s) were generated as predictions, using the fitted parameters. The longer delay leads to smaller difference in amplitudes (ΔH_1_ < ΔH_2_). The WPI, but not the WP model accounts for these experimental features. The WP model fails to capture the heterogeneity of the data, resulting in a wider spread. This also reduced the number of successful fits from a total of 81 (successful WPI model fits) to only 58 (for WP).

The WP and WPI models were fit as before, with PIP3 data used as a surrogate for the stimulus to Rac. The parameters found from the fitting procedure were used to solve the WP and WPI models and resulted in predictions shown in the center and right columns of **Figure VIII**. The WPI (but not the WP) model was able to account for several features of the experimental observations, namely the relative height of the first and second peaks (ΔH_1_, ΔH_2_, see right panels), and the “undershoot” of Rac activity (which fell below its unstimulated level) between and after the stimuli. The WP model failed to reproduce these observations.

### Results 2D: The WPI model responds to stimulation rate

We investigated the WPI model’s response to the *rate* of stimulation. Building on results of **Figure VII**, we asked how the WPI model responds to a gradual “ramp-up” stimulus, in place of the previous ON-OFF pulses. The predictions of the WPI model and the corresponding experimental results, are shown in **Figure IX**. As shown in **Figure IX**, (bottom row), the WPI model predicts a controlled rise in Rac activity without the overshoot seen in the stimulus pulses (**Figure IX**, top row). However, the model also predicts a slight increase in Rac activity above pre-stimulus levels, contrary to experimental observations, where Rac levels remain close to baseline.

**Figure IX:**
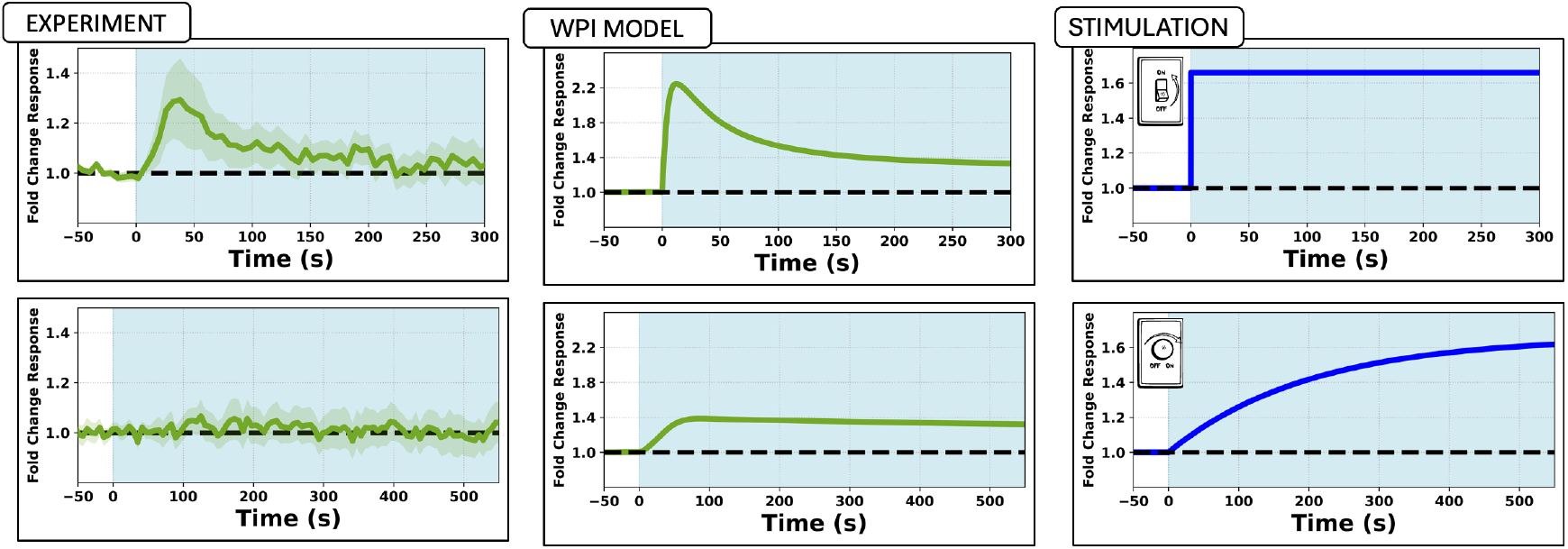
Responses to ON versus ramp stimuli. The WPI model distinguishes between abrupt and gradual stimuli **(Center)**. A step-function stimulus **(Top)** triggers an overshoot in Rac activity, followed by adaptation, in agreement with experiments **(Left)** of Town & Weiner (2023). A gradual stimulus **(Bottom)** leads to a mild Rac increase above baseline (in contrast with the baseline data) with no overshoot. The findings suggest that the Rac inhibitor filters stimuli, reacting to rapid, rather than gradual changes.

These findings suggest that the WPI model responds more strongly to rapid stimuli than to gradual changes or absolute levels. Thus, it appears that the inhibitory component serves as a filter, enabling the cell to differentiate effectively between rapid and gradual changes in its environment.

### Result 3A: WPI responds more strongly to sudden stimuli than to gradual stimuli

We next sought to determine how the temporal response of the WPI model to stimulation rates translates into a spatio-temporal response to localized stimuli along a cell’s edge. Consequently, we returned to the spatial (PDE) version of the WPI model and examined its response to spatially localized stimuli that were either instantaneous (pulse) or gradual (ramp) in time.

As shown in **Figure X**, we initiated the model with a constant stimulus at −90° on the 1D wrap-around cell edge domain. At *t = 100s*, the second stimulus, introduced at 90° was either a pulse (top) or a gradually increasing stimulus (bottom), implying high and low rates of change, respectively. Despite both sites eventually receiving equal levels of stimulation, the responses (shown as kymographs in **Figure X**) were distinct.

**Figure X:**
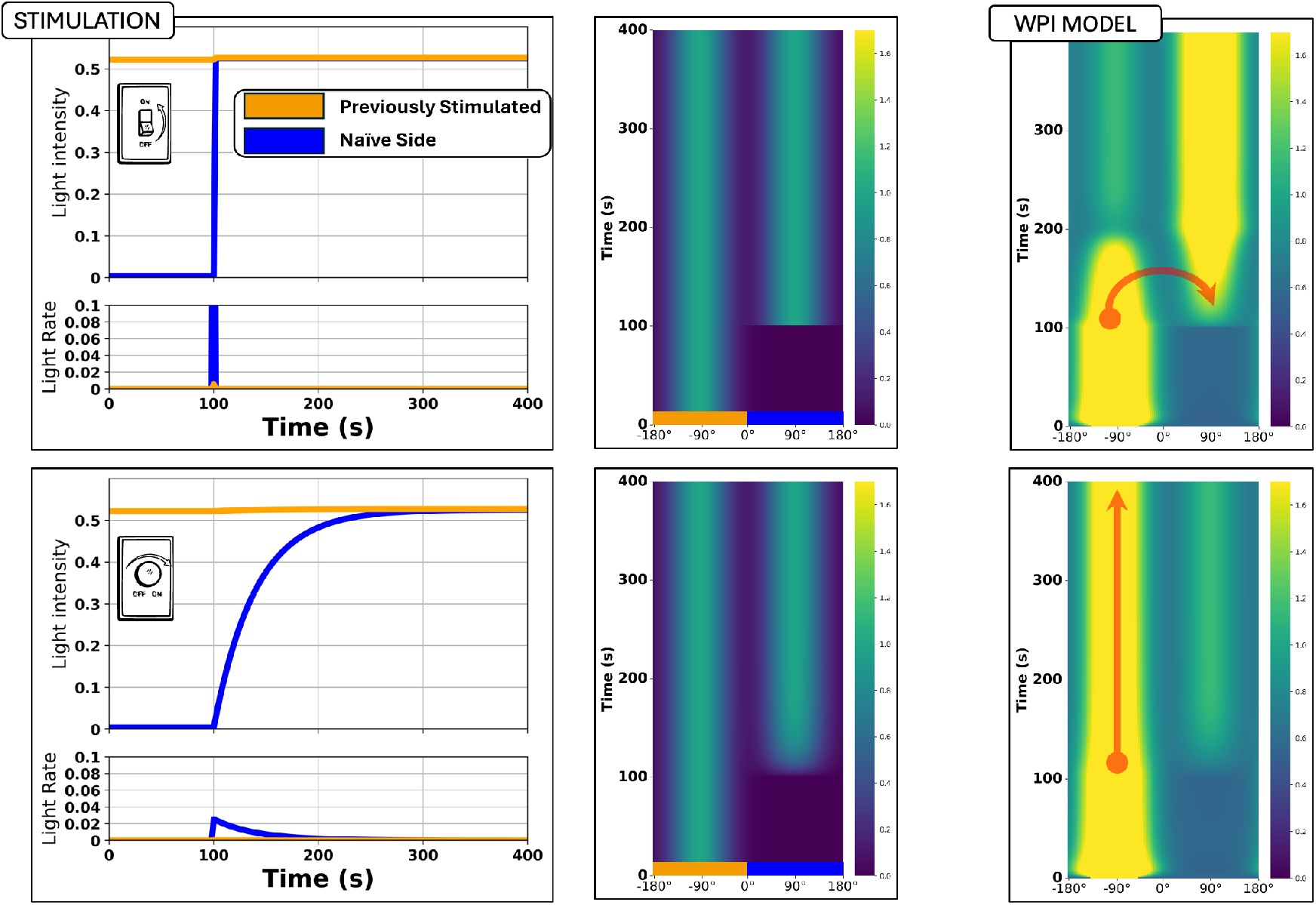
Spatial responses to abrupt versus gradual stimuli. The WPI model differentiates between abrupt and gradual stimuli in a spatial context. Model cells were initially exposed to a constant stimulus on one side, followed by the application of a competing stimulus on the opposite side. When the competing stimulus was introduced as a step-function **(top)**, it redistributed the active Rac to its location. Conversely, when the competing stimulus was applied gradually **(bottom)**, the location of the active zone of Rac remained unchanged.

In response to the abrupt second stimulus (**Figure X**, top row), Rac activity rapidly shifted from its previous site (−90°) to the newly stimulated site. In contrast, in response to the gradual input (**Figure X**, bottom row), the original site retained its Rac activity, abrogating the formation of the competing new Rac zone at 90°. These results reinforce the idea that the WPI model responds more strongly at sites on the cell that experience more rapid stimuli, over those regions that experience sustained, graduate stimuli, i.e., that the rate of change of input is the key signal.

In contrast to the WP model where the intensity of the stimulus and/or its time duration is the signal to repolarize (Buttenschön & Edelstein-Keshet, 2022), in the WPI model, the Rac inhibitor effectively filters out gradual stimuli, preserving existing zones of Rac activity unless a sufficiently strong and/or rapid perturbation occurs. This mechanism could be biologically relevant in guiding directional responses, allowing cells to ignore weak or slowly changing signals while rapidly adjusting to new, more urgent cues.

### Results 3B: After guidance, cells with WPI continue rotating

We asked how the WP and WPI models play out in directing motility of a model cell. To visualize cell shape and motility, we simulated cells using the cellular Potts model in Morpheus, as previously described. We implemented the WP and WPI signaling models on the edge of such simulated cells (See SI) using the software Morpheus.

We first considered polarized cells exposed to simple local stimuli that lead to cell turning. These results are shown in Supplementary **Figure S3**. Briefly, we found that both WP and WPI models can account for the experimentally observed response. In both cases, the model cell turns, as expected, after stimulation. In both cases, the light stimuli were fixed in space, allowing us to redirect the cell to a fixed direction.

In response to a more complex stimulus designed to guide the cell into rotation, we observe a more intriguing result. Initially, we stimulate the cell on its left side (90_0_ counterclockwise from its movement direction), causing the cell to rotate (shown in **Figure XI**). When the cell’s front was facing north, we stopped the stimulation. As previously shown in **Figure VIII**, Rac activation responds faster to the stimulation than the inhibitor. Spatially, this means that the inhibitor lags behind the Rac zone, effectively pushing Rac forward and sustaining its motion.

**Figure XI:**
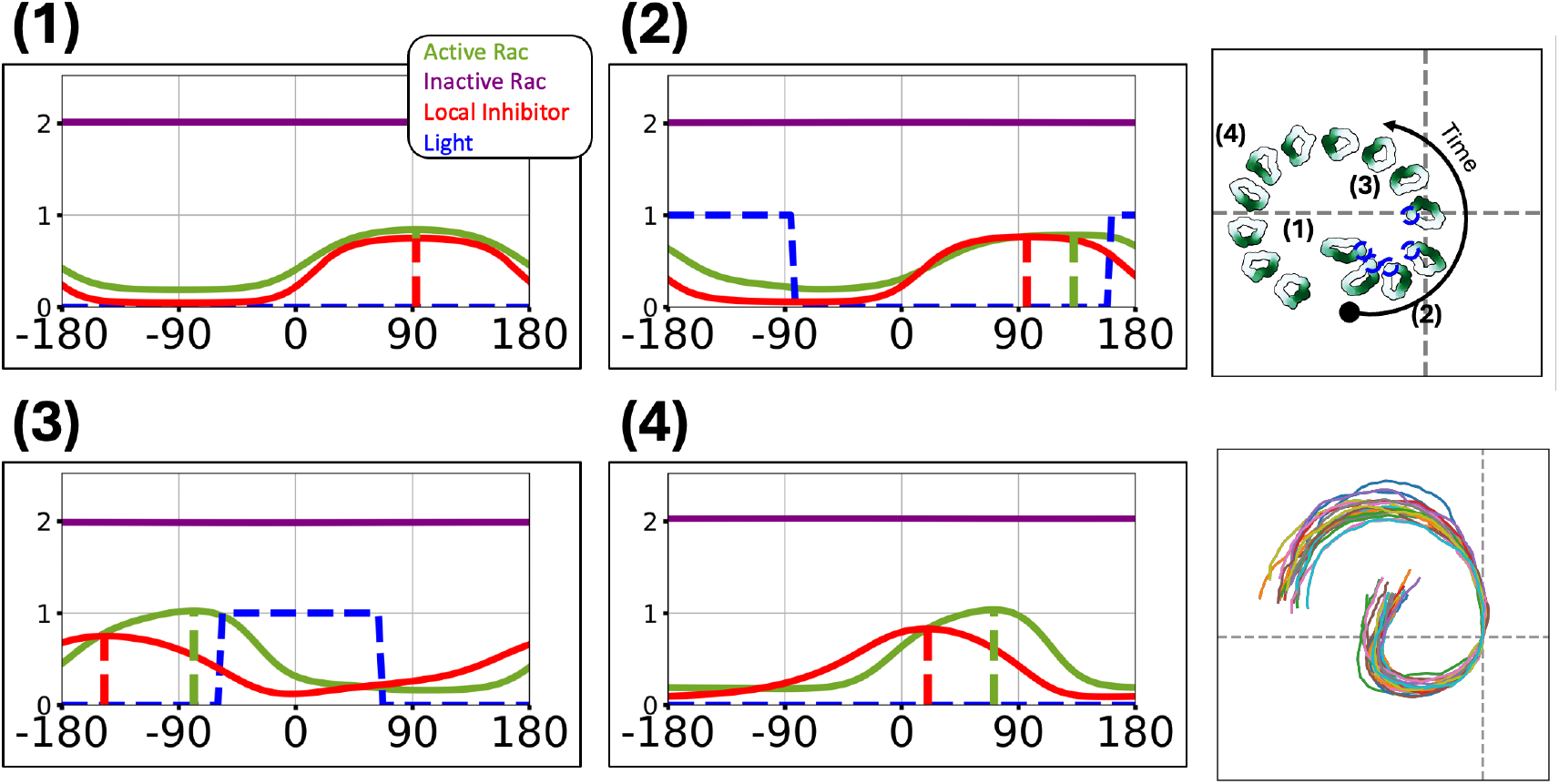
Guiding the cell to rotate. The WPI model is shown, implemented on the “unwrapped” 1D cell edge (left four panels) and on a CPM simulation in Morpheus **(Right)**. **(Left)** Four time points showing spatial profiles of active and inactive Rac (green, purple, respectively), the Rac inhibitor (red), and the light stimulus (blue). The peaks of active Rac and inhibitor are indicated (dashed lines). **(Top right)** Snapshots of the motile cell moving in 2D at corresponding time points. Active Rac is shown in green shading, and blue circles indicate light stimulus. The inhibitor trails behind Rac, sustaining a persistent rotation of the cell. Time points denote **(1)** period before stimulation, **(2**,**3)** periods of direct stimulation, **(4)** periods after stimulation. Video shown in Supplemental Material: **Movie 3**

### Results 4A: Parameters obtained by fitting the models

Table 1 summarizes the best-fit values of model parameters obtained by the fitting algorithm. We used only the double-pulse experiment shown in the lower left panel of **Figure VIII** (for Δt_1_=120s) to fit parameters and then simulated the other cases with the set of parameters so obtained. Details about the method are given in the SI/Appendix.

**Table 1:** Model parameters fit to experimental data. Fitted parameter values for the WP (residuals_2_ = 4183.57, n = 58) and WPI (residuals_2_ = 615.19, n =81) models. The stated values and ranges represent a 95% confidence level given the distribution of fitted parameters. The Hill coefficient for Rac autocatalysis was kept fixed at n=2. Time is in seconds (s). Data and models are scaled so that active Rac is normalized (given as fold-multiples of its basal prestimulus level). To capture the heterogeneity, data for many cells were pooled together. Each cell has its own PIP3 level (used as proxy for input stimulus) and its own normalized Rac activity level. The inhibitor is in arbitrary units [AU]. For details about the fitting routine, see the Supplemental Information.

| Parameters | WP | WPI |
| --- | --- | --- |
| $T$ (Total Rac) [AU] | 9.78±0.70 | 7.31 ± 0.28 |
| $k_0$ (Basal Rac activation rate) [1/s] | 0.035±0.006 | 0.028 ± 0.003 |
| $s$ (Stimulus-induced Rac activation rate) [1/s] | 0.12±0.03 | 0.059 ± 0.006 |
| $\gamma$ (Autocatalytic Rac activation rate) [1/s] | 1.31±0.10 | 1.38 ± 0.084 |
| $K$ (Hill-function parameter for Autocatalytic) [AU] | 2.75±0.19 | 2.93 ± 0.16 |
| $\delta$ (Basal Rac inactivation rate) [1/s] | 2.70±0.11 | 0.721 ± 0.027 |
| $k_1$ (Inhibitor induced Rac inactivation) | n/a | 0.372 ± 0.026 |
| $\alpha$ (Rac-dependent inhibitor production rate) [1/s] | n/a | 6.8x10<sup>-3</sup>± 4.0x10<sup>-4</sup> |
| $\delta_h$ (Inhibitor decay rate) [1/s] | n/a | 4.0x10<sup>-3</sup>± 6.0 x10<sup>-4</sup> |

**Table 2:** Model selection. Akaike Information Criterion (AIC) model comparison using the best-fit parameters for the temporal data corresponding to the 120s gap. The AIC values were calculated using observations from all time-points (30s, 60s, 120s, 240s, and 480s), highlighting that the WPI model provides a significantly better fit to the data than the WP model. See Supplementary Information for further details.

| Model | AIC | ΔAIC | AIC Weight | RSS | Samples (n) |
| --- | --- | --- | --- | --- | --- |
| WPI | 938.6 | 0 | ~1.00 | 3707.5 | 425 |
| WP | 1160.3 | 221.7 | ~0.00 | 6335.8 | 425 |

**Table 3:** Relative rates of diffusion fit to experimental data. Setting the diffusion coefficient of active Rac to D_u_=1 (in arbitrary units), we scaled all other diffusion coefficients relative to D_u_. Fitting was performed across all spatial stimulation protocols (n = 166), using the average Rac profile within each protocol. As expected, inactive Rac diffuses faster than active Rac. We also found that the inhibitor diffuses most slowly. See SI for correspondence to unit-carrying values and for further details.

| Parameters | WPI | WPI-PIP3 |
| --- | --- | --- |
| $D_u$ (Active Rac diffusion rate) | 1 | 1 |
| $D_v$ (Inactive Rac diffusion rate) | 3.37 | 6.40 |
| $D_H$ (Inhibitor diffusion rate) | 0.14 | 0.47 |

We find several interesting aspects of the parameter values so obtained. (1) The basal Rac activation rate (*k*_*0*_) is small, and much smaller than the elevated rate when Rac is autoactivating itself (*k*_*0*_ << γ). Indeed, the basal rate is around 3% of the elevated activation rate (*k*_*0*_ ~0.027 γ). This is required to assure a low resting-level of Rac activity that can get significantly elevated by positive feedback. (2) The basal rate of Rac inactivation (δ) is considerably higher (by 3.75-fold) in the WP model that has no inhibitor. This makes sense, since in the WPI model, much of the work of damping out Rac activity is accomplished by the putative inhibitor. (3) The level of total Rac (*T*) is larger in the WP model than in the WPI (34%). This difference can be understood by realizing that for long periods of stimulation, WP holds a steady level of activation, while WPI rises and then falls (following adaptation). Hence when fitting the models to data, WP was forced to balance between the peak and the long-term Rac activity, resulting in fits that favor higher values of the Rac activation rates. On the other hand, WPI can reach high peaks immediately after stimulation, followed by lower long-term activity thanks to the slow inhibitor time dynamics. (4) In the WPI model, the rate of decay of the inhibitor (δ_h_) is significantly lower (<1%) than the rate of decay of Rac. This implies that the inhibitor persists for much longer (half-life = ln(2)/0.0040 ~ 173s), compared to Rac (half-life=ln(2)/0.721 ~ 1s). (5) This is the first time that experimental data is being used to fit parameters to the WP model (since the Mori et al 2008 paper), and results of the fits are highly consistent with ball-park estimates made in that purely theoretical model. In particular, it was noted there and in Hughes et al (2024) that a nonzero basal rate of Rac activation (*k*_*0*_ > 0) is theoretically essential for polarity by the WP mechanism to exist, but that it should be very small compared to the autocatalytic rate of Rac activation due to the positive feedback of Rac (*k*_*0*_ << γ). The fits to the WP model, confirm this theory, even while the WPI model is seen to be a better fit to the neutrophil stimuli experiments.

It would be interesting to compare the parameter values with others from the literature, but the number of previous model fits to data is quite limited. Lockley et al (2014), (and Lockley PhD thesis (2015) Appendix A) fit three polarity models (Meinhardt 1999, Otsuji et al 2007, and Levchenko & Iglesias 2002) to F-actin data from Dictyostelium cells repolarizing in response to reversal of a shear flow. These models do not specifically identify molecular components, but both Meinhardt’s and Otsuji have an “active” variable (*A, U*, respectively). Otsuji has the pair *U, V* for active and inactive forms of the protein, with mass conservation, as does the Mori et al (2008) WP model, though other details differ significantly. The leading rate of activation and decay of *U* in Otsuji et al (a_1_~0.38/s), and the inactivation rate of *A* in Meinhardt (r_a_~0.23/s) are on the same order of magnitude as the rate γ for WP-Rac activation and the basal WPI-Rac inactivation rate δ.

Other papers that adopt variants of the WP model, e.g., the Wang et al (2017) cancer cell study, also use a speculative set of parameters, or at best, fit one or two parameter values to the model.

Rates of diffusion (in μm^2^s^−1^) that were fit for the various previous model components differ vastly. For the Otsuji model, the ratio of rates of diffusion of *V* and *U* was taken to be 10^6^, much higher than needed by the WP model. For the Meinhardt model, the ratio D_A_/D_C_, was on the order of 0.5. Since these differ greatly from ours and from one another, it is not possible to draw meaningful comparisons for such parameter values.

For our simulations, we fitted the diffusion rates and obtained a ratio of 6.4 between active and inactive Rac. This value aligns well with parameter estimates and choices in previous studies (Mori et al 2008, Hughes et al 2024) for the WP model.

### Results 4B: Selection of best fit time-dependent model

The time-dependent WPI model has three parameters more than the WP model, as seen in Table 1. We asked whether the WPI model is a better parsimonious fit to the data despite this increase in degrees of freedom. To address this question, we used the Akaike Information Criterion (AIC) which rewards a low sum of squared errors (SSE) and penalizes the number of parameters in comparing distinct models fit to the same dataset. The details of how we computed the AIC score are given in the supplemental material.

### Result 4C: Fitting spatial Rac polarization data to motile cells

So far, we have only considered temporal data and non-spatial models for Rac activity in immobilized latrunculin-treated cells. Next, we examine spatial models using data from motile cell polarization which, for the cells we study here, depends on F-actin activity to maintain distinct front and back regions. This aligns with the Wave-Pinning (WP) model, where polarization results from different diffusion rates of active and inactive Rac. Inactive Rac diffuses quickly, staying nearly uniform across the cell, while active Rac forms a polar pattern. As polarization occurs, inactive Rac is depleted from the cytoplasm, preventing new polarization sites. Prior studies (Mori et al., 2008; Hughes et al., 2024) noted that the WP model is rather sensitive to parameters. In our hands, the WP model indeed proved difficult to fit reliably to spatial data. We therefore proceeded with the WPI and WPI-PIP3 models, keeping all parameter values that we fit from the time-dependent model.

Fitting the spatial data requires the full spatio-temporal model to account for the spatial spread of both Rac activity and Rac inhibitor. Hence, rates of diffusion for these variables also have to be estimated or fit. We report relative diffusion ratios normalized to the active Rac diffusion rate (set to 1), physical units and scaling details are provided in the SI. Fitting was performed across multiple spatial stimulation protocols. Further details can be found in the SI/Appendix.

Similar to the previous fitting, our results aligned with theoretical expectations: (1) The diffusion rate of active Rac is found to be small, while inactive Rac has a much larger diffusion coefficient (approximately 6.4-fold higher). (2) The hypothesized inhibitor has a diffusion rate lower to that of active Rac, reinforcing its local nature.

### Results 5A: WPI does not account for reversal experiment

The core puzzle addressed in Town & Weiner (2023), (see **Figure I**) is the counterintuitive reversal of cell rotation following a shift from local to global stimulation. In experiments, when a cell is locally stimulated on its left side, it rotates counterclockwise (as shown in **Figure XI**).

However, upon switching to a static global stimulus in the experiments, most cells reverse direction and rotate clockwise. Based on arguments in Town & Weiner (2023), we Initially hypothesized that the WPI model would suffice to capture this behaviour based on two key findings: (1) **Result 3C** shows that the WPI model responds to stimulation *rates*, suggesting that abrupt stimuli should lead to significant changes in Rac activity. (2) **Result 3D** demonstrates that rapid spatially localized stimuli would allow a new Rac zone to gain advantage and grow, despite continued stimulation of a pre-existing zone.

Given these results, it stands to reason that an abrupt transition from local to global stimulus should leave the original (left) Rac zone unchanged, while promoting a new Rac zone in the (uninhibited) right region of the cell. We argued that, in principle, this sudden change would cause the new Rac zone to outcompete the existing zone, relocating the “front” of the cell, and driving the reversal of motion seen in the experimental data.

To test this, we implemented the local-to-global stimulation protocol as in experiments. To ensure a continuous turning, we stimulated the cell at a site 90° counterclockwise from its front (previously denoted as the “left side of the cell”). As soon as the cell pointed north, the local stimulus was abruptly switched to a global one. This r*apid* switch should have introduced a large, sudden rate of Rac activation at 90° clockwise of the existing front, potentially triggering reversal. To reflect biological variability across cells, we sample from the pool of fitted parameter sets when simulating multiple trajectories, allowing us to explore population-level responses and heterogeneity in behaviours such as polarization or reversal.

Surprisingly, the WPI model failed to reproduce the reversal. Instead, it persists in counterclockwise rotation, in contrast to the experimentally observed switch. **Figure XII** illustrates this result, showing the light stimulus, its rate of change, and the resulting active Rac around the cell’s edge. Peaks of active Rac and inhibitor (indicated with green and red lines, respectively), show how the inhibitor lags behind Rac.

**Figure XII:**
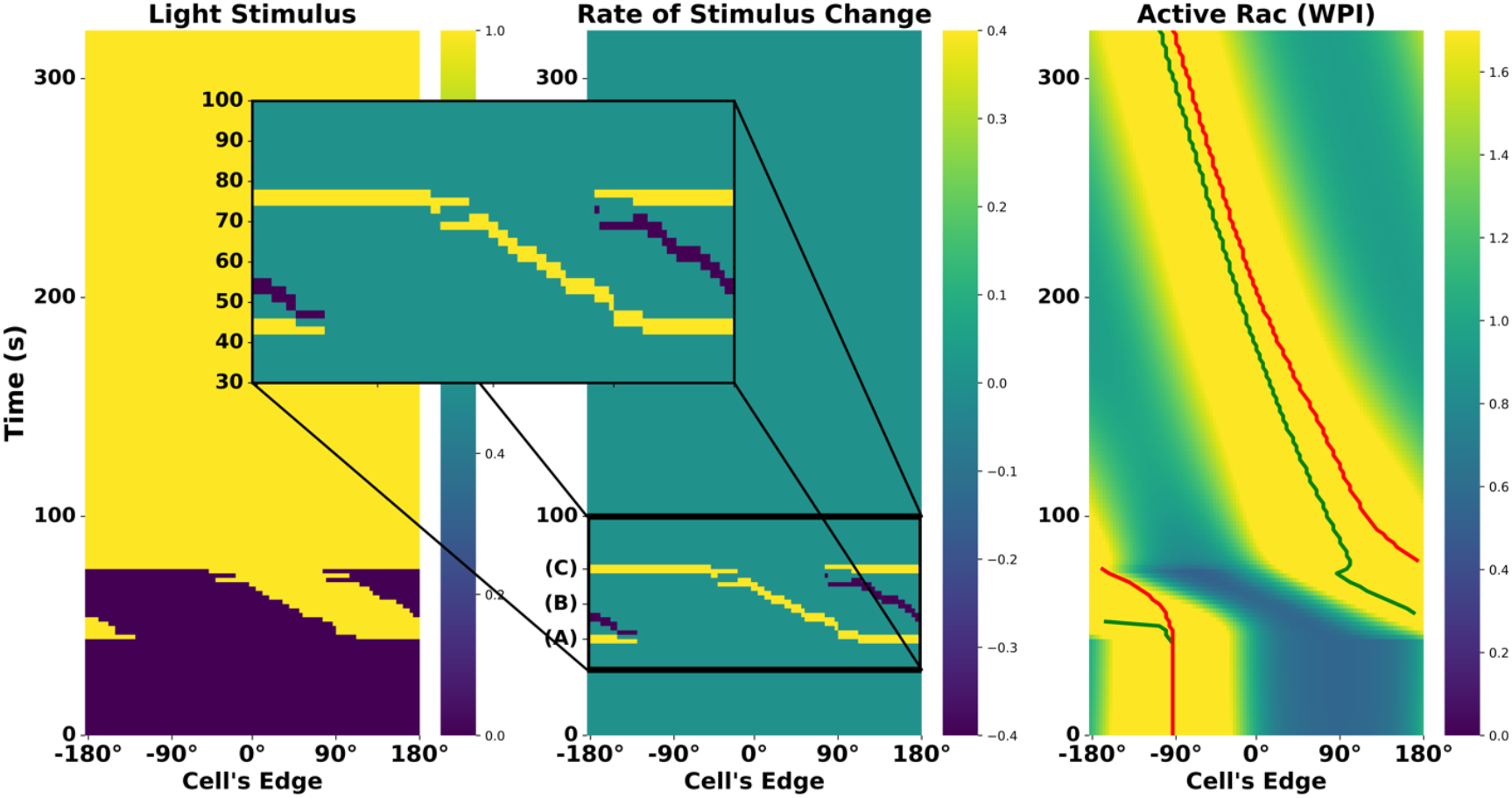
WPI model response to the local-to-global reversal experiment. Left to right: kymographs of light stimulus, its rate of change, and active Rac. The zoom-in highlights three key moments: **(A)** onset of local stimulation, **(B)** transient phase during rotation, and **(C)** switch to global stimulation. The WPI model fails to reverse Rac polarity following global stimulus, despite a sharp rate of change at the back of the active zone. Green and red curves in the right panel track the peak positions of Rac and the inhibitor, respectively

Key insights from these results suggest that while the rate of change in stimulus plays a crucial role (as seen in **Results 3D**), it is not the only factor governing Rac redistribution. From **Results 3E**, we learned that Rac maintains persistent rotation due to the timescale differences between its activation and inhibitor dynamics (**Table 1**). This insight confirms that the inhibitor lags behind Rac with a strong enough influence to prevent reversal.

Although the rate of change at the right of the Rac zone is indeed large and rapid, it occurs over a very short period of time. The inhibitor, on the other hand, exerts an effect over a longer period, overcoming the transient stimulus shift and preserving the original rotation.

These findings reveal a fundamental limitation of the WPI model: while it captures stimulus-dependent Rac redistribution, it fails to account for persistent inhibitory lag that appear to be crucial for explaining reversal.

### Results 5B: Lag due to PIP3 explains reversal

Having established that the WPI model alone fails to capture the experimentally observed reversal (**Figure XII**), we sought a minimal extension that could explain this behaviour. Our previous results (**Figure X, Figure IX**) demonstrated that Rac redistribution is sensitive to the rate of stimulus change, but in **Figure XII**, we saw that this effect alone was not sufficient to drive reversal. Instead, Rac persisted in its original rotation.

The data in Town & Weiner (2023) includes the time course of PIP3 upstream of Rac activation. Including this component in our model easily recapitulates the reversal track. Having a parametrized set of models allows us to ask what features of the expanded WPI-PIP3 model account for the reversal and under what conditions. In the SI, we describe a sequence of in silico experiments that help to pinpoint the necessary conditions and how interactions between the components leads to reversal. Briefly, we show that (1) PIP3 adds a lag that maintains the influence of the local stimulus for a short while during global stimulation. (2) If PIP3 dynamics are too fast or too slow, there is no reversal. (3) Indeed, the cell’s reversal depends on a delicate balance between all parameters. When we sample parameter sets from our fitted distributions (mimicking biological heterogeneity), we observe cases where reversal does not occur, in agreement with experimental results. We briefly summarize these observations in **Figure XIII, XIV** and show details in Supplementary **Figures S2-S3**.

**Figure XIII:**
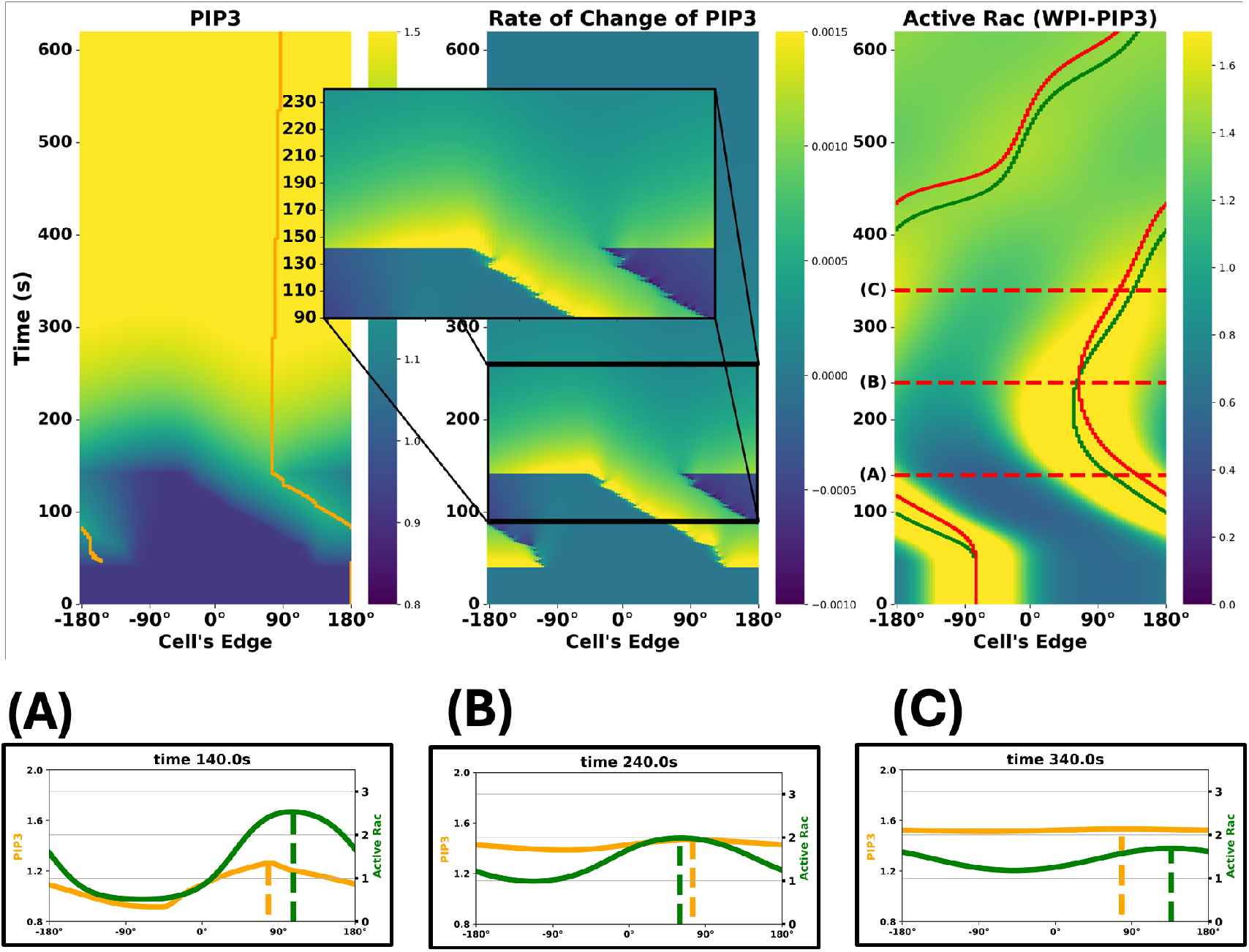
The WPI-PIP3 model accounts for the reversal experiment. **(Top)** Kymographs of: PIP3 concentration **(Left)**, PIP3 rate of change **(Center)**, and active Rac **(Right)**, with orange, green, and red curves tracking the peaks of PIP3, Rac, and inhibitor, respectively. Time points (on kymograph, right and bottom row): **(A)** Global stimulus turned on, **(B)** Rac and PIP3 peaks switch positions, (just before the inhibitor catches up, not shown), **(C)** reverse rotation initiated. The spatial profiles of PIP3 (orange) and active Rac (green) are shown at corresponding time points **(bottom row)**. Dashed lines denote peaks of PIP3 and Active Rac.

**Figure XIV:**
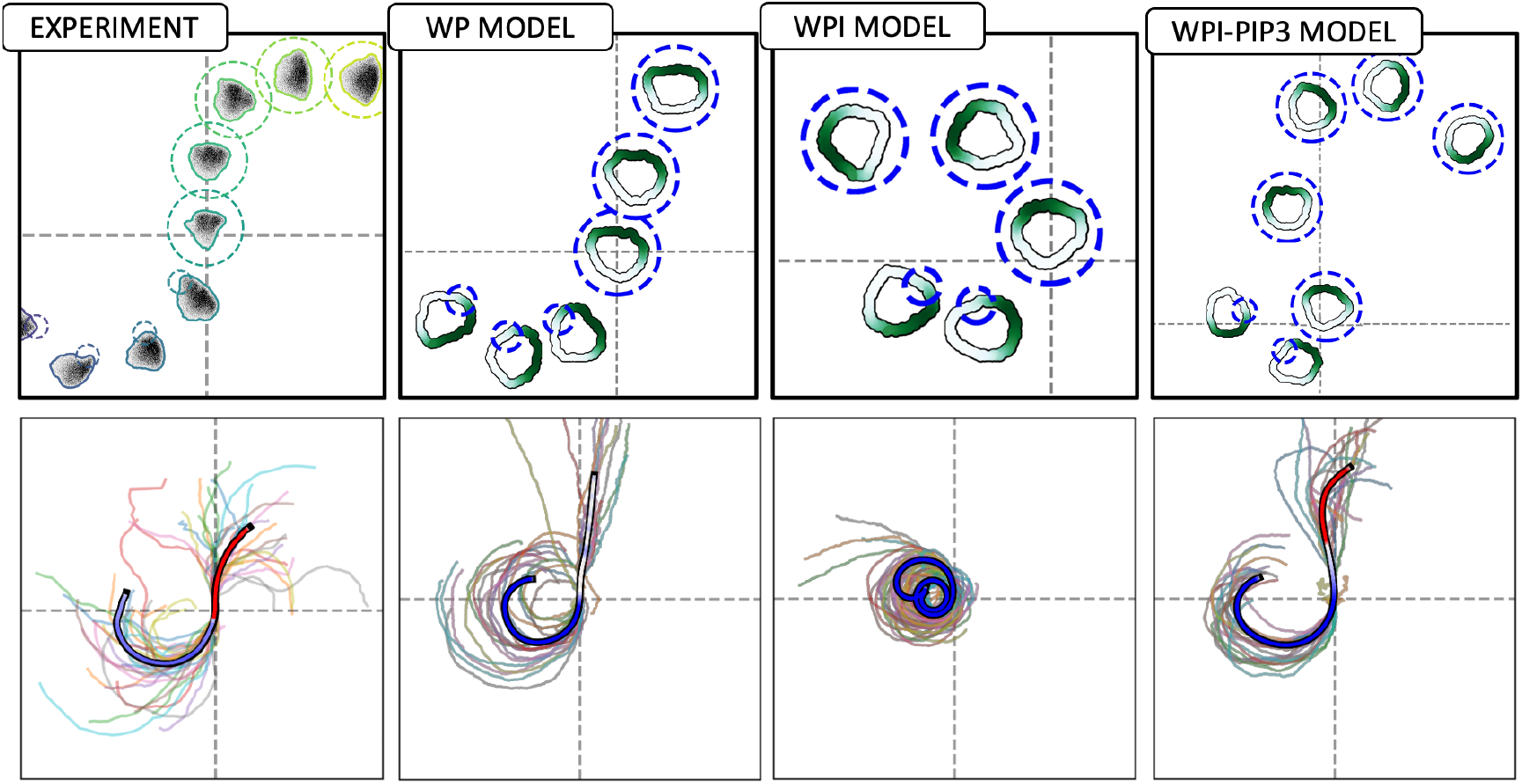
The WPI-PIP3 (but not WP or WPI) model accounts for reversal trajectories. A comparison of cell shapes, active Rac distributions along the cell edge (green) and various cell trajectories in the experiment (left) and in cell simulations predicted by the three models (L to R: WP, WPI, and WPI-PIP3) with heterogenous parameters. In the lower panels, all individual cell tracks are shown (25 for the experiment, 20 for each of the models), with the average trajectory overlaid and colour-coded by rotation direction: blue for counterclockwise (CCW), red for clockwise (CW), and white for no rotation. In all models, the cell is first locally stimulated on its left from t = 100s to t = 300s, triggering counterclockwise rotation. At t = 300s, global stimulus is turned on. WP predicts straight migration or loss of polarity and stalling; WPI continues its initial rotation; Only WPI-PIP3 reverses direction, matching experimental results. Video shown in Supplemental Material: **Movie 4**

### Results 6: Model cell responses to dynamic and noisy gradients

Having extracted meaningful parameter values from all experimental data, we can now test the responses of the model cell to virtual environments beyond those already tested experimentally, such as noisy and dynamic chemical gradients. To do so, we assumed that each point along the cell edge transduces the local attractant chemical concentration into an elevated rate of local Rac activation. (This replaces the on-off 0-1 optogenetic light stimulus term previously used). The gradient of chemical results in a gradient of Rac activation, determining the Rac zone, the cell front, and hence, the direction and rate of migration.

We simulated a scenario where cells are initially placed in a northward pointing chemical gradient. At *t = 80*, the chemical field is briefly reset to uniform, followed by gradient reversal at *t = 120*. As shown in **Figure XV**, all three models (WP, WPI, and WPI-PIP3) initially polarize and undergo chemotactic migration up the gradient. Based on our fitted parameters, we found that the WP model predicts delayed polarization onset compared with WPI and WPI-PIP3, particularly in regions of low chemoattractant concentration. Upon gradient reversal, only the inhibitor models (WPI and WPI-PIP3) successfully reoriented. WP remained fixed in its original direction. WPI cells showed a slower reversal, as shown by their short reversal paths. In contrast, WPI-PIP3 cells reversed more rapidly, producing longer reversed trajectories.

**Figure XV:**
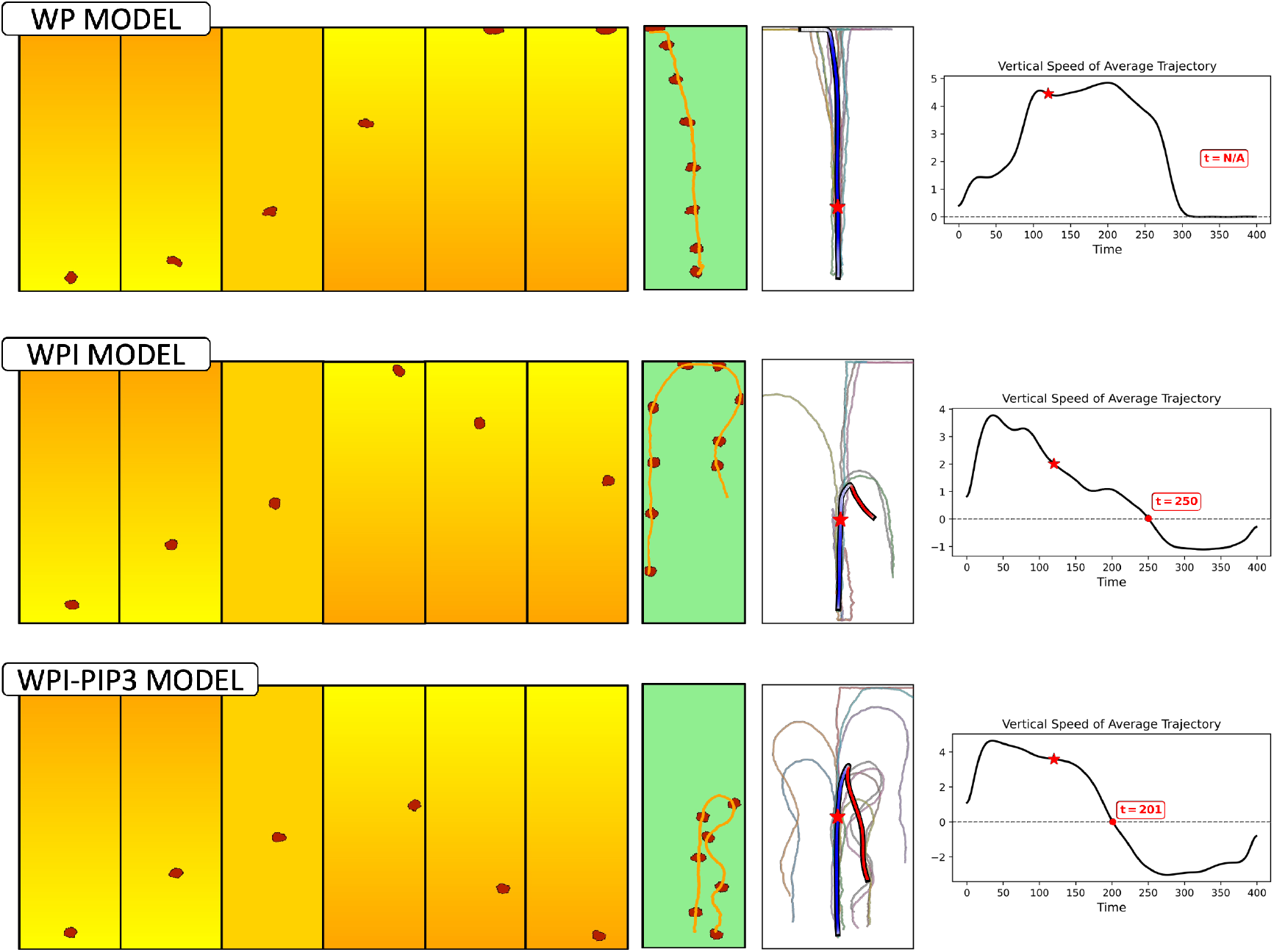
Model cell responses to dynamic chemotactic gradients. Predicted sensitivity of the three types of model cells to dynamic chemical gradients. Left six panels: snapshots of cell positions (red shapes) and chemical gradient from low (yellow) to high (orange) concentrations. In a noise-free gradient, eventually all cells align with the initial gradient. WP (top) polarizes very slowly and fails to reorient after gradient reversal. In contrast, WPI and WPI-PIP3 (middle and bottom) polarize faster and successfully reverse direction after the gradient flips. Green and white panels: single representative cell trajectory, followed by a sample of several. The blue-red coloured track indicates the average path taken by the cells. The colour changes from blue (Northwards direction) to red (Southwards direction), with a red star marking the gradient reversal timepoint. Right panel: average vertical velocity vs time (with the same red star); the red dot indicates the cell reorientation time, (undefined for the WP model). Notably, WPI-PIP3 reorients faster (t = 201) than WPI (t = 250). The sudden drop in WP vertical speed is due to the cell reaching the top boundary of the domain, not active reorientation. Video shown in Supplemental Material: **Movie 5**

The differences in the models can be explained as follows. First, the inhibitor accumulates at the front and destabilizes the Rac zone there. This destabilization facilitates the replacement of the existing front, making reversal more likely. Second, the faster reorientation in the WPI-PIP3 model arises from the PIP3 dynamic adaptation to the changing gradient. After the gradient reverses, PIP3 redistributes from a previously high level, decreasing across the cell, but more slowly in the direction of the new gradient. This creates an early asymmetry that facilitates a faster response.

Additional tests assessing model performance in noisy gradients are presented as rose plots in the Supplementary Information. These reinforce the same conclusion: the WPI-PIP3 model shows the most robust alignment across a variety of environmental conditions.

## Discussion

In this study, we developed and analysed a series of basic, biologically grounded models for Rac activation and polarization in neutrophils. Our main contribution is to provide a modelling framework that validates a key experimental hypothesis proposed by Town and Weiner (2023): that a local Rac inhibitor is required to explain responses to optogenetic stimuli. Through a systematic investigation of several model variants, we demonstrate and explained the roles of a Rac inhibitor and of PIP3 in accounting for responses to various stimulation protocols, most importantly, the reversal experiment. These results show that combining a local inhibitory mechanism with a slow-adapting intermediate (PIP3) provides a robust and minimal mechanism for flexible polarity responses in motile cells.

The behaviours observed in our simulations suggest several biological implications. Most notably, we find that the ability to reorient in response to changing cues, such as a switch from local to global stimulation or a gradient reversal, requires both a local inhibitor and a dynamic “filtering” component such as PIP3. The inhibitor plays a primary role in enabling redirection by destabilizing the current front. PIP3 further enhances reorientation by smoothing stimulus transitions and preserving a directional bias after abrupt changes. This extended memory-like behaviour helps the system avoid erratic responses, particularly in noisy environments. Together, the inhibitor and PIP3 work in concert, to provide a basic biologically plausible mechanism for robust polarity control in migrating cells.

Although this work has been focused on neutrophil cellular migration, this model could be generalized to other amoeboid cells such as *Dictyostelium*, where the dynamics of Rac have been studied in the past (Park et al., 2004; Šoštar et al., 2024). Several modelling studies have previously explored the dynamics of cell polarity and migration using reaction-diffusion systems coupled to deformable domains. A previous mode closest to ours is Neilson et al (2011) where a putative pseudopod activator and two inhibitors (local and global) were assumed in reaction-diffusion PDEs on a deforming closed curve representing the edge of a motile 2D cell. Their simulation method has the advantage of including the effects of edge curvature and “membrane tension’’ on the RD system. Our CPM is better at depicting fluctuating 2D cell shapes, while identifying specific model components and fitting to experimental data. Other works, such as Marée et al. (2006) introduced a multiscale model of keratocyte motility that combines GTPase signalling with actin-driven protrusions and mechanical feedback. More recently, Liu et al. (2021) extended the WP framework to include source-sink mechanisms and actin feedback, simulating various patterning regimes and cell shape behaviours using a custom Cellular Potts Model implementation. In a parallel modelling framework, Camley et al. (2017) employed a phase-field approach to couple shape dynamics with molecular dynamics.

The rate of diffusion of the inhibitor was found to be smaller than that of active Rac. This suggests that it, too, is membrane-bound, and possibly of larger molecular weight. The diffusion ratio of 1.0:0.47 (active Rac : inhibitor) implies that the inhibitor could have MW roughly (1.0/0.47)_3_ =9.6 times that of Rac. Since typical small GTPases are around 21 kDa, this implies the inhibitor MW is on the order of 200 kDa. This is in line with some of the larger GAPs, such as ARHGAP5 (172 kDa) and MYO9B (243 kDa). Alternately, the inhibitor could have protein-binding domains that lead to formation of complexes around that size, also a feature known in GAPs.

Our parameter fitting results also imply that the inhibitor is produced or (activated) at a very low rate by Rac, while also having a very slow decay (or inactivation) rate relative to Rac kinetics. These results may eventually lead to further identification of the likely molecular candidate(s) playing the role of the Rac inhibitor.

While previous studies provide important insights into self-organization and shape change, our work differs in three keyways. First, we test specific hypotheses about the molecular circuit underlying polarity reversal, using both time-dependent simulations and cell motility simulations to account for experimental observations. Second, rather than relying solely on hand-tuned parameters, our model is quantitatively calibrated to experimental data, enabling us to extract parameter distributions across cells and thereby account for biological heterogeneity. Third, our use of the Morpheus platform makes the full CPM model openly accessible and easily reproducible, lowering the barrier for further exploration and validation.

While our model reproduces key aspects of polarity reversal and directional adaptation, it simplifies several biological processes. In particular, the dynamics are simulated in a one-dimensional periodic domain, which removes a degree of freedom for movements of molecules. We also do not include mechanical feedback from membrane tension with has been shown to act as a global inhibitor of Rac dynamics (Houk A et al., 2012). Furthermore, the model assumes that polarity dictates movement, without feedback from cell deformation. The intracellular dynamics are also simplified, focusing on a minimal Rac-inhibitor-PIP3 circuit and omitting additional regulators such as Rho or Cdc42. These simplifications allow us to isolate and test for minimal requirements for polarity reversal but also point to natural directions for future model refinement.

## Acknowledgments

This work was supported by National Institutes of Health grant GM118167 (ODW), and by a Natural Sciences and Engineering Research Council (NSERC, Canada) Discovery Grant (LEK).

## References

Burnham KP, Anderson DR, SpringerLINK eBooks - English/International Collection (Archive), Springer Books, Ebook Central, SpringerLink (Online service). Model Selection and Multimodel Inference: A Practical Information-Theoretic Approach. 2nd ed. New York: Springer; 2002;2003.

Buttenschön A, Edelstein-Keshet L. Cell repolarization: A bifurcation study of spatio-temporal perturbations of polar cells. Bulletin of Mathematical Biology. 2022 Oct;84(10):114.

Camley BA, Zhao Y, Li B, Levine H, Rappel WJ. Crawling and turning in a minimal reaction-diffusion cell motility model: Coupling cell shape and biochemistry. Physical Review E. 2017 Jan;95(1):012401.

Chang H, Levchenko A. Adaptive molecular networks controlling chemotactic migration: dynamic inputs and selection of the network architecture. Philosophical Transactions of the Royal Society B: Biological Sciences. 2013 Nov 5;368(1629):20130117.

Graner F, Glazier JA. Simulation of biological cell sorting using a two-dimensional extended Potts model. Physical review letters. 1992 Sep 28;69(13):2013.

Guntas G, Hallett RA, Zimmerman SP, Williams T, Yumerefendi H, Bear JE, Kuhlman B. Engineering an improved light-induced dimer (iLID) for controlling the localization and activity of signaling proteins. Proceedings of the National Academy of Sciences. 2015 Jan 6;112(1):112–7.

Hadjitheodorou A, Bell GR, Ellett F, Shastry S, Irimia D, Collins SR, Theriot JA. Directional reorientation of migrating neutrophils is limited by suppression of receptor input signaling at the cell rear through myosin II activity. Nature Communications. 2021 Nov 16;12(1):6619.

Houk A, Jilkine A, Mejean C, et al. Membrane Tension Maintains Cell Polarity by Confining Signals to the Leading Edge during Neutrophil Migration. Cell. 2012;148:175–188.

Inoue T, Meyer T. Synthetic activation of endogenous PI3K and Rac identifies an AND-gate switch for cell polarization and migration. PloS one. 2008 Aug 27;3(8):e3068.

Jilkine A, Edelstein-Keshet L. A comparison of mathematical models for polarization of single eukaryotic cells in response to guided cues. PLoS computational biology. 2011 Apr 28;7(4):e1001121.

Levchenko A, Iglesias PA. Models of eukaryotic gradient sensing: application to chemotaxis of amoebae and neutrophils. Biophysical journal. 2002 Jan 1;82(1):50–63.

Lin B, Holmes WR, Wang CJ, Ueno T, Harwell A, Edelstein-Keshet L, Inoue T, Levchenko A. Synthetic spatially graded Rac activation drives cell polarization and movement. Proceedings of the National Academy of Sciences. 2012 Dec 26;109(52):E3668–77.

Liu Y, Rens EG, Edelstein-Keshet L. Spots, stripes, and spiral waves in models for static and motile cells: GTPase patterns in cells. Journal of Mathematical Biology. 2021 Mar;82:1–38.

Lockley R, Ladds G, Bretschneider T. Image based validation of dynamical models for cell reorientation. Cytometry Part A. 2015 Jun;87(6):471–80.

Lockley R. Image-based modelling of cell reorientation (Doctoral dissertation, University of Warwick). 2015.

Marée AF, Jilkine A, Dawes A, Grieneisen VA, Edelstein-Keshet L. Polarization and movement of keratocytes: a multiscale modelling approach. Bulletin of mathematical biology. 2006 Jul;68:1169–211.

Marée AF, Grieneisen VA, Edelstein-Keshet L. How cells integrate complex stimuli: the effect of feedback from phosphoinositides and cell shape on cell polarization and motility. PLoS computational biology. 2012 Mar 1;8(3):e1002402.

Meinhardt H. Orientation of chemotactic cells and growth cones: models and mechanisms. Journal of cell science. 1999 Sep 1;112(17):2867–74.

Mori Y, Jilkine A, Edelstein-Keshet L. Wave-pinning and cell polarity from a bistable reaction-diffusion system. Biophysical journal. 2008 May 1;94(9):3684–97.

Neilson MP, Mackenzie JA, Webb SD, Insall RH. Modeling cell movement and chemotaxis using pseudopod-based feedback. SIAM Journal on Scientific Computing. 2011;33(3):1035–57.

Neilson MP, Veltman DM, van Haastert PJ, Webb SD, Mackenzie JA, Insall RH. Chemotaxis: a feedback-based computational model robustly predicts multiple aspects of real cell behaviour. PLoS biology. 2011 May 17;9(5):e1000618.

Niculescu I, Textor J, De Boer RJ. Crawling and gliding: a computational model for shape-driven cell migration. PLoS computational biology. 2015 Oct 21;11(10):e1004280.

Otsuji M, Ishihara S, Co C, Kaibuchi K, Mochizuki A, Kuroda S. A mass conserved reaction– diffusion system captures properties of cell polarity. PLoS computational biology. 2007 Jun;3(6):e108.

Park KC, Rivero F, Meili R, Lee S, Apone F, Firtel RA. Rac regulation of chemotaxis and morphogenesis in Dictyostelium. EMBO J. 2004 Oct 7;23(21):4177–4189. doi: 10.1038/sj.emboj.7600368

Price, Kenneth. Differential Evolution: A Practical Approach to Global Optimization. Edited by Jouni A. Lampinen, and Rainer M. Storn. Springer Nature, Berlin, Heidelberg, 2006;2005;, doi:10.1007/3-540-31306-0.

Shi C, Huang CH, Devreotes PN, Iglesias PA. Interaction of motility, directional sensing, and polarity modules recreates the behaviors of chemotaxing cells. PLoS computational biology. 2013 Jul 4;9(7):e1003122.

Šoštar M, Marinović M, Filić V, Pavin N, Weber I. Oscillatory dynamics of Rac1 activity in Dictyostelium discoideum amoebae. PLoS Comput Biol. 2024;20(12):e1012025. doi: 10.1371/journal.pcbi.1012025

Starruß J, De Back W, Brusch L, Deutsch A. Morpheus: a user-friendly modeling environment for multiscale and multicellular systems biology. Bioinformatics. 2014 May 1;30(9):1331–2.

Toettcher JE, Gong D, Lim WA, Weiner OD. Light-based feedback for controlling intracellular signaling dynamics. Nature methods. 2011 Oct;8(10):837–9.

Town JP, Weiner OD. Local negative feedback of Rac activity at the leading edge underlies a pilot pseudopod-like program for amoeboid cell guidance. PLoS Biology. 2023 Sep 25;21(9):e3002307.

Vanderlei B, Feng JJ, Edelstein-Keshet L. A computational model of cell polarization and motility coupling mechanics and biochemistry. Multiscale Modeling & Simulation. 2011 Oct 1;9(4):1420–43.

Wang W, Tao K, Wang J, Yang G, Ouyang Q, Wang Y, Zhang L, Liu F. Exploring the inhibitory effect of membrane tension on cell polarization. PLoS computational biology. 2017 Jan 30;13(1):e1005354.

Zmurchok C, Collette J, Rajagopal V, Holmes WR. Membrane Tension Can Enhance Adaptation to Maintain Polarity of Migrating Cells. Biophysical journal. 2020;119:1617–1629.

