## Supplemental Information for "Investigating local negative feedback of Rac activity by mathematical models and cell motility simulations"

#### 1 Modelling Methods

##### 1.1 Model geometry

We use two frames of reference, the lab coordinates, and a coordinate system along the cell edge. In lab coordinates, $0^o$ is taken to be in the "North" direction, for convenience, and most simulations start with cells polarized in that direction. We also consider a cell-based coordinate system to describe positions along the cell edge. "Front" then denotes the location of maximal Rac activity on the edge, rear is at the opposite pole of the cell, and right or left sides are at $\pm\pi/2$ radians from the front location.

The cell frame of reference moves relative to lab coordinates. As the cell moves, stimuli in [1] were computer-controlled to be fixed relative to this inherent cell framework. (Hence, for example, "stimulating on the left" always implies a stimulus at $-\pi/2$ from the "front".)

In the models described below, we use $-180 \leq x < 180$ to denote the lab frame of reference, analogous to a lab "compass". Hence, as the cell reorients due to stimuli, the Rac zone (as well as the stimulus, and all other variables) will appear to "shift" relative to this frame of reference.

##### 1.2 Reaction-Diffusion Equations

To investigate the spatiotemporal dynamics of Rac, we started with a basic reaction-diffusion (RD) model for Rac cycling between active and inactive states proposed by Mori et al. (2008) [2], to account for cell polarization. This model, denoted the "wave-pinning (WP) model", was implemented on a one-dimensional (1D) periodic ("wrap-around") domain representing the cell edge.

Let $u(x,t)$ and $v(x,t)$ denote the levels of active and inactive Rac at position $x$ along the cell perimeter ($\Omega$) at time $t$. The governing equations for the original WP model are:

$$\frac{\partial u}{\partial t} = v\left(k_0 + \gamma\frac{u^n}{K^n + u^n}\right) - \delta u + D_u\frac{\partial^2 u}{\partial x^2}, \quad (1a)$$

$$\frac{\partial v}{\partial t} = -v\left(k_0 + \gamma\frac{u^n}{K^n + u^n}\right) + \delta u + D_v\frac{\partial^2 v}{\partial x^2}. \quad (1b)$$

These equations have the property that the total amount of Rac, $T$, is constant in the domain.

$$T = \int_\Omega (u + v)\, dx, \quad (2)$$

where the model parameters are: $k_0$ (basal Rac activation rate), $\gamma$ (maximal Rac autoactivation rate), $K$ (Rac level inducing 50% maximal autoactivation), $n$ (Hill coefficient, $n = 2$), $\delta$ (basal Rac inactivation rate), $D_u$ and $D_v$ (diffusion rates of active and inactive Rac, respectively), and $T$ (total Rac, conserved over the timescale of interest).

Mori et al. [2] showed that, under appropriate conditions, the model (1) can account for spontaneous symmetry breaking consistent with formation of a polar pattern of Rac. Conditions for such spontaneous polarization include distinct rates of diffusion, $D_u \ll D_v$. This disparity in diffusion rates is a natural consequence of GTPase biology: the active form is bound to the membrane, a much more viscous environment than the cytosol in which the inactive form diffuses. The distinct diffusion rates between $u$ and $v$ create an effective global interaction where inactive Rac rapidly redistributes across the domain, acting as a pool of Rac whose depletion halts the spread of the Rac zone. For a polar pattern to form, there are also conditions on $T$. (See [2]).

The RD dynamics are implemented on the 1D cell edge, as Morpheus does not support deforming 2D or 3D domains. This 1D simplification is similar to prior works by [3] and [4].

##### 1.3 The Cellular Potts Model (CPM) Simulations

To simulate cell motility driven by Rac dynamics, we implemented the model equations (1) on the edge of a 2D deformable simulated cell in Morpheus (Starruß et al., [5]). Morpheus is an open source multiscale modelling platform for simulating single and collective cell behavior, based on the Cellular Potts Model (CPM).

CPM cells are represented by a connected set of pixels with a fluctuating edge that can randomly protrude or retract at each point. An energetic cost function (the "Hamiltonian") prevents the total area and/or perimeter of the CPM cell from deviating away from a user-specified target area ($A_0$) and perimeter ($P_0$). These constraints are weighted by strengths $\lambda_a, \lambda_p$ set by the user. In general, we set $\lambda_a = 1, \lambda_p = 1$, and pick $P_0$ as a fold-multiple, $\phi$, of the perimeter of a circle with area $A_0$. The constant $\phi$ denoted the "aspherity", describes the deviation of a cell shape from that of a perfect circle. In general,

$$\text{Aspherity} = \frac{P}{2\sqrt{\pi A}},$$

where $P$ is the perimeter and $A$ the area of the cell. For a perfect circle, this ratio equals 1. Values greater than 1 indicate increasingly elongated or irregular cells shapes. (We found $\phi \approx 1.4 - 1.46$ to be consistent with shapes of neutrophils.) The Morpheus software allows all such parameters to be easily specified. For fuller details about the Cellular Potts Model, see [6].

While previous custom-built CPM code represented PDEs for GTPase activity inside 2D deforming CPM cells [7, 8], in open-source software such as Morpheus, this type of simulation is not yet available. However, the Morpheus plugin `MembraneProperty` allows users to solve reaction-diffusion equations such as (1) in a periodic 1D domain representing the "cell edge"[1].

To couple the Rac activity to protrusion at points along the cell's edge, we used the Morpheus plugin `StarConvex`; this plugin allows us to favours protrusive pixel fluctuations at locations along the cell-edge with high Rac activity (high $u$). This idea is consistent with localized nucleation of F-actin downstream of active Rac, and hence, protrusion associated with high Rac activity zones. Furthermore, to represent implicit "persistence", with gradual (rather than instantaneous) cytoskeletal turnover (while not explicitly tracking F-actin in the model), we used the Morpheus

[1] In actual fact, Morpheus solves such problems on a circular domain with the same area as the CPM cell, and then "projects" the solution values along radii from the cell's centroid to its perimeter.

`Protrusion` plugin. This plugin implements the "Act" idea of Niculescu et al. [9] [2], enabling the cell to sustain and amplify the protrusions caused by `StarConvex`.

The RD system is implemented in Morpheus using the `DiffEqn` plugin, which allows direct numerical integration of PDEs on domains such as the cell edge. The edge distribution of active Rac is solved on a discretized 1D lattice, using the sequential operator splitting method. The diffusion is solved using the central difference method, while the reaction is solved with the Euler method time-stepping.

#### 1.4 Building up the Model and Modelling the Stimulus:

##### 1.4.1 The stimulated WP model

In optogenetic experiments, Rac activation is controlled with high spatial and temporal precision through light input. To model this, we added a stimulus term, $S(x,t)$, to the rate of activation of Rac in the original WP model, as in previous work [10]:

$$\frac{\partial u}{\partial t} = v\left(k_0 + \alpha S(x,t) + \gamma\frac{u^n}{K^n + u^n}\right) - \delta u + D_u\frac{\partial^2 u}{\partial x^2}, \tag{3a}$$

$$\frac{\partial v}{\partial t} = -v\left(k_0 + \alpha S(x,t) + \gamma\frac{u^n}{K^n + u^n}\right) + \delta u + D_v\frac{\partial^2 v}{\partial x^2}. \tag{3b}$$

For optogenetic stimuli, we took $S(x,t) = 0, 1$. Local (global) stimuli were implemented as $S = 1$ on a small interval along the cell edge ($S = 1$ uniformly). $\alpha$ controls the effective stimulus-induced rate of Rac activation. In modelling the response of cells to a reversing "chemical gradients" (Figure XV and Figures S4-S5), $S(x,t)$ was taken to represent the chemical concentration at the position of the cell membrane, mapped from a global extracellular field , as in [10].

The light-stimulated WP model fails to explain key features of the experimental data, both time-dependent responses and cell trajectories following a switch from local to global stimulation.

##### 1.4.2 The WPI model

Following the hypothesis proposed by Town and Weiner [1], we added a Rac inhibitor, $h$ to the model system. We assumed that inhibitor is produced downstream of active Rac, and amplifies Rac inactivation. Denote the inhibitor by $h(x,t)$. For simplicity, assume that inhibitor is produced at a rate proportional to active Rac ($u(x,t)$), and degrades with simple first-order kinetics. Then the RD equation for the inhibitor is

$$\frac{\partial h}{\partial t} = k_h u - \delta_h h + D_h\frac{\partial^2 h}{\partial x^2}, \tag{4a}$$

where the inhibitor parameters are: $k_h$ (inhibitor production rate), $\delta_h$ (degradation rate), and $D_h$ (inhibitor diffusion rate). The extended model includes a modification of (1), assuming that Rac

[2] As in [9], newly added pixels are assigned a fixed "actin level" (ACT) that decays over time, creating a memory of recent protrusive activity. The CPM Hamiltonian is biased to favour copy attempts from high-activity pixels to neighbouring low-activity pixels, mimicking actin-driven edge protrusion.

inactivation increases linearly with the level of the inhibitor, $h$. The new equations for the WPI version of the model become:

$$\frac{\partial u}{\partial t} = v\left(k_0 + \alpha S(x,t) + \gamma \frac{u^n}{K^n + u^n}\right) - (\delta + k_1 h)\, u + D_u \frac{\partial^2 u}{\partial x^2}, \tag{4b}$$

$$\frac{\partial v}{\partial t} = -v\left(k_0 + \alpha S(x,t) + \gamma \frac{u^n}{K^n + u^n}\right) + (\delta + k_1 h)\, u + D_v \frac{\partial^2 v}{\partial x^2}. \tag{4c}$$

Here, the inhibitory term $k_1 h$ introduces a negative feedback loop. The coefficient $k_1$ controls the strength of this inhibition. This feedback enables the system to suppress Rac activity after some stimulation is given, becoming an adaptive system [11], a characteristic not observed in the original WP model.

The WPI model successfully captures adaptive features observed in experiments (Figures VIII, IX). WPI distinguishes between slow and rapid stimulation (Figure X) and predicts persistent cell turning (Figure XI). However, it fails to reproduce the reverse turning cell paths observed when the stimulus is switched from local to global (Results 5A).

###### 1.4.3 The WPI-PIP3 model

In the Main Text, we briefly explain the role of PIP3 and why its inclusion in the model is important (Results 5B) PIP3, a signalling lipid, is produced from PIP2 by the kinase PI3K (activated optogenetically). PIP3 subsequently activates Rac.

We denote the level of PIP3 by $p(x,t)$, and assume it is produced from a basal pool of PIP2 in response to light stimulus:

$$\frac{\partial p}{\partial t} = (T_p - p)\left(k_p + \beta S\left(x,t\right)\right) - \delta_p p + D_p \frac{\partial^2 p}{\partial x^2}, \tag{5a}$$

where the model parameters are: $T_p$ (total amount of PIP3 and PIP2 in the cell), $k_p$ (basal PIP3 production rate), $\beta$ (light stimulus strength coefficient), $\delta_p$ (basal PIP3 degradation rate), and $D_p$ (PIP3 diffusion rate). The Rac activation term is then modified by replacing the direct light term $\alpha S(x,t)$ with a dynamic PIP3 input. The new system of equations becomes (5a), (4a) and:

$$\frac{\partial u}{\partial t} = v\left(k_0 + \alpha p(x,t) + \gamma \frac{u^n}{K^n + u^n}\right) - (\delta + k_1 h)\, u + D_u \frac{\partial^2 u}{\partial x^2}, \tag{5b}$$

$$\frac{\partial v}{\partial t} = -v\left(k_0 + \alpha p(x,t) + \gamma \frac{u^n}{K^n + u^n}\right) + (\delta + k_1 h)\, u + D_v \frac{\partial^2 u}{\partial x^2}. \tag{5c}$$

This system forms the WPI-PIP3 model. The key idea is that PIP3 introduces a time delay between light input and Rac activation due to its own dynamics. This extension introduces no new feedback loops but delays and spreads the effect of the light signal in space and time. The resulting behaviour of the model aligns more closely with experimental results, particularly in reproducing polarity reversal after global stimulation.

##### 1.5 Mechanism of Rotation Reversal in the WPI-PIP3 Model

Because PIP3 production is driven by light, it accumulates within the Rac zone during local stimulation, forming a nonuniform pattern with a peak. When the stimulus switches to a global stimulus, this localized PIP3 peak does not immediately dissipate. Instead, it persists briefly, creating a transient spatial bias.

Immediately after we switch to the global stimulus, Rac continues its counterclockwise rotation, driven by the delayed peak of the inhibitor, which sustains the previous motion. However, as Rac progresses, it leaves behind the PIP3 peak that formed during local stimulation. This lingering PIP3 peak locally enhances Rac activation and effectively pulls Rac back toward it, acting as a temporary “anchor point” or attractor. As a result, Rac slows down and eventually reverses direction. By the time PIP3 becomes uniform across the cell, Rac has already reoriented and the inhibitor, now trailing the shifted Rac, helps maintain the reversed rotation.

We tested this mechanism in two ways:

- **Constant inhibitor:** To isolate the role of adaptive feedback, we removed the dynamic inhibitor and set its effect to a constant value. This essentially reduces the system back to a WP-like regime, with Rac inactivation fixed across space and time. During local stimulation, Rac slowly shifts towards the stimulus site. However, upon switching to global stimulation, memory of the previous stimulation is entirely lost. The Rac zone then stalls at its last position and maintains a stable front. This mirrors behaviour seen in the standard WP model and confirms that a dynamic inhibitory response is necessary for directional change (Figure S1)

- **PIP3 Rate variation:** To test the sensitivity of the model to PIP3 dynamics, we altered the PIP3 timescale in the full WPi-PIP3 model (with dynamic inhibitor). Results in Figure S2demonstrate that for fast PIP3 dynamics, PIP3 quickly becomes uniform across the domain. Once global stimulation is applied. Without any lingering asymmetry, Rac experiences a spatially uniform stimulus from PIP3 and continues rotation under the influence of the inhibitor, effectively behaving like the WPI model. On the other hand, for very slow PIP3 dynamics, the PIP3 peak remains localized for too long, acting as a fixed attractor. Rac is unable to fully pass or disengage from it, resulting in oscillatory motion around the PIP3 peak rather than a clean reversal.

These tests demonstrate that reversal depends on a fine balance between a delayed, localized inhibitor and a transient, asymmetric PIP3 signal. If either component is missing or mistimed, the system either fails to reverse (WPI-like) or becomes trapped, losing polarity (WP-like).

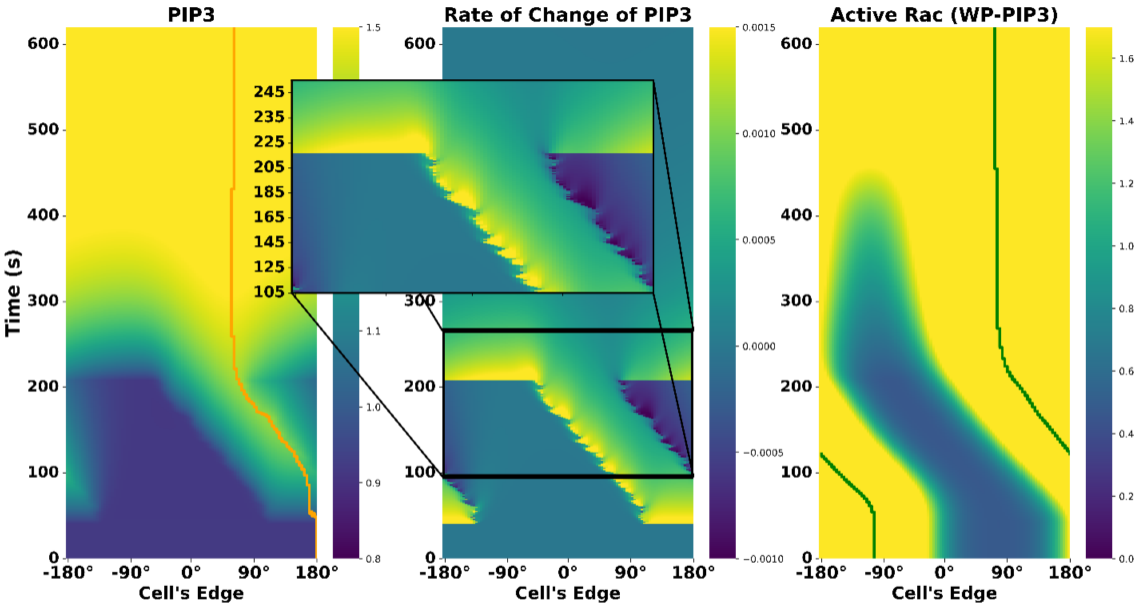


Figure S1: Simulations of the WP model with PIP3 and inhibitor held constant, showing response for local-to-global stimulation. Left to right: kymographs of PIP3, its rate of change, and active Rac. Upon global stimulation, the Rac zone remains fixed in place until polarization is lost due to sustained stimulation.

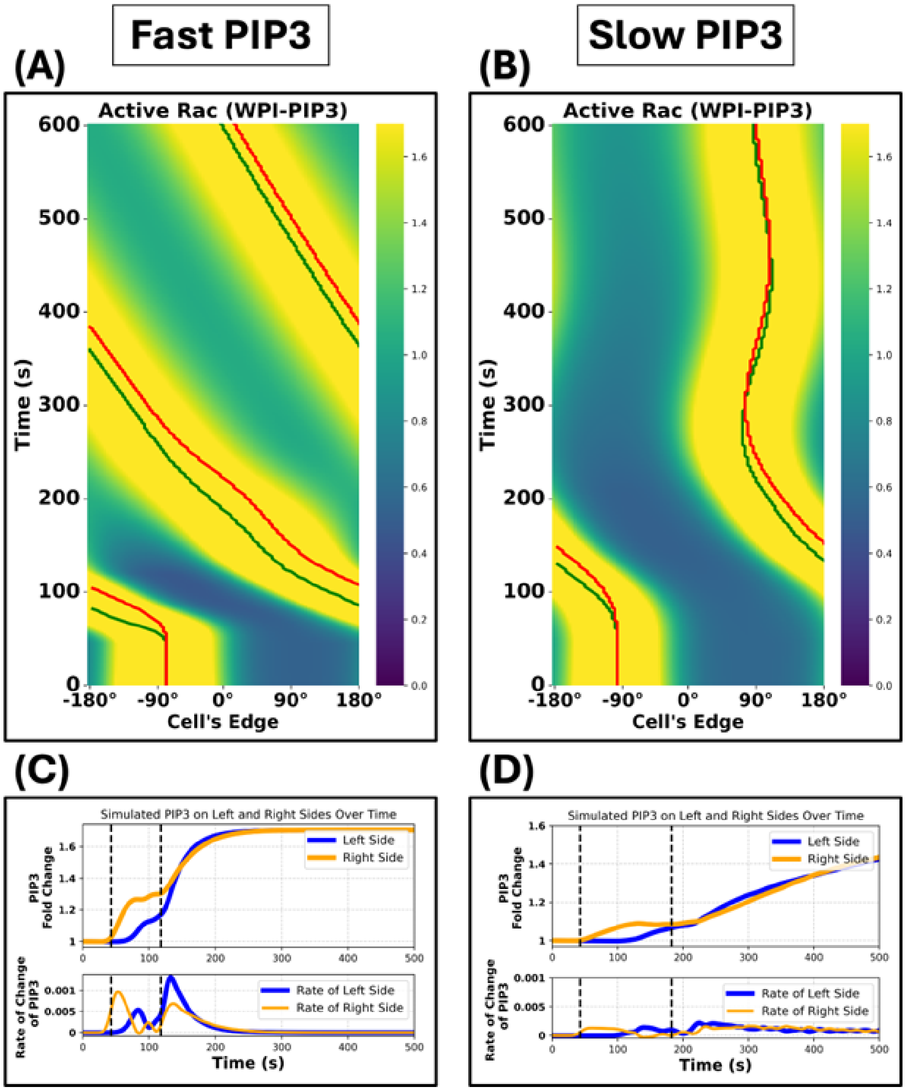


Figure S2: Effect of PIP3 dynamics on Rac response. (Top) Kymographs of active Rac for fast (A) and slow (B) PIP3 dynamics, obtained by globally scaling the PIP3 equation with a timescale perameter. Green and red curves track the peaks of Rac and inhibitor, respectively. (Bottom) Time-dynamics of PIP3 (fold change, top graph) and its rate of change (bottom graph) on the left and right sides of the cell. Dashed vertical lines indicate the onset of local stimulation (first line) and the switch to global stimulation (second line). In the fast PIP3 case, PIP3 becomes uniform across the domain (equal levels on left and right side) before Rac can respond to any residual asymmetry, resulting in continued rotation. In the slow PIP3 case, the PIP3 peak persist too long, anchoring Rac and leading to oscillatory drift rather than full reversal. Only intermediate PIP3 dynamics (not shown) enable transient directional bias, producing successful reversal.

#### 2 Fitting Parameters:

To accurately map our model simulations to real cellular dimensions, we first extracted the characteristic size of neutrophil-like HL-60 cells from the imaging data provided in [1]. The dataset in [1] includes a conversion factor from image pixel to physical units µm, which we used to quantify the perimeter, area, and asphericity of the cells.

| Geometric measure | Value ± Standard Deviation |
|---|---|
| Perimeter | 76.0 ± 14.12 µm |
| Major Diameter | 25.2 ± 3.88 µm |
| Minor Diameter | 8.82 ± 2.26 µm |
| Area | 222.2 ± 75.95 µm$^2$ |
| Aspherity | 1.46 ± 0.048 |

Table S1: Geometric characteristics of motile HL-60 cells in the reversal experiment from Town & Weiner (2023). Values represent the mean ± standard deviation for perimeter, area, diameters and aspherity.

To increase spatial fidelity, we tripled the resolution of the experimental pixel-to-µm ratio in our CPM simulations, and rescaled diffusion rates to preserve consistent dynamics.

##### 2.1 Time-Dependent Reaction Parameters

To fit the temporal dynamics of Rac activity in latrunculin-treated cells, we used a least-squares minimization procedure to identify parameter sets that best reproduce the experimental Rac responses shown in Figures VII and VIII. These datasets track fold-changes in Rac activity over time following a double-pulse protocol: cells were stimulated globally for 180 seconds, followed by a rest phase (no stimulation) of 120 seconds, and then stimulated again for another 180 seconds. Because latrunculin inhibits F-actin polymerization and immobilizes the cells, spatial effects and feedback from shape change were absent, enabling us to simplify the model to its ODE form with only the reaction kinetics. In these experiments, light stimulation activates PI3K, leading to the production of PIP3, which in turn promotes Rac activity. The observed variability in PIP3 responses provided a range of inputs, enabling us to capture some of the heterogeneity in Rac dynamics across cells by fitting parameters to each individual cell.

For each cell $j$, we minimized the cost function

$$C_j = \sum_{i=1}^{N} \left(R_{ij}^{\text{exp}} - R_{ij}^{\text{sim}}\right)^2 ,$$

where $R_{ij}^{\text{exp}}$ is the experimental fold-change of Rac in cell $j$ at timepoint $t_i$, and $R_{ij}^{\text{sim}}$ is the corresponding simulation prediction. Here, $N$ denotes the number of interpolated timepoints ($N \approx 85{,}000$). Rac activity was normalized to a baseline value of 1 for each cell using its pre-stimulus level.

We used the Differential Evolution algorithm from the `scipy.optimize` package to perform the global optimization. DE is a stochastic population-based method that iteratively improves parameter estimates using mutation, crossover, and selection operations (Price, K et al., [12]).

The DE optimizer was initialized using Latin Hypercube Sampling to ensure broad coverage of parameter space. Each ODE was solved using the `solve_ivp` function from the `scipy.integrate` package.

Before each simulation, the system was run to steady state to determine pre-stimulus levels, allowing us to create a baseline for computing fold change. To fit the double-pulse dataset, the PIP3 fold-change data were interpolated from experimental measurements and treated as a known time-dependent input to Rac. This allows us to isolate Rac and inhibitor dynamics from uncertainties in upstream signalling.

The parameters obtained through this fitting procedure are listed in Main text Table 1. In total, we obtained 58 successful fits for the WP model and 87 for the WPI model. (The WP model is known to be rather sensitive.) These fits produced parameter distributions from which we later sampled in order to preserve biological heterogeneity in our simulated cells.

#### 2.2 AIC Model Comparison

To assess whether the WPI model provides a better fit than the WP model while accounting for its addtional parameters, we computed the Akaike Information Criterion (AIC) for both models using their best-fit residuals and number of parameters. AIC score is defined as:

$$\mathrm{AIC} = 2k + n \ln(\mathrm{RSS}/n),$$

where $k$ is the number of fitted parameters, $n$ the number of data points, and RSS is the residual sum of squares.

We applied this calculation to the temporal fits from all double-pulse stimulation experiments. For this dataset, both models were evaluated using the same 425 individual cells, using their PIP3 data directly as a known input function for $p(x, t)$ in the model. The WPI model had three additional parameters over the WP model, due to the inhibitor dynamics.

The resulting AIC values are shown in Table 2. The difference in $\Delta$AIC was substantial, indiciating strong evidence in favour of the WPI model.

#### 2.3 Fitting Diffusion Parameters from Spatial Data

To estimate the relative rates of diffusion for active and inactive Rac, Rac-inhibitor, and PIP3, we simulated the full spatio-temporal WPI-PIP3 model. The reaction-diffusion equations were discretized in space using second-order central differences in a periodic 1D domain, with $M = 100$ grid points. This yields a tridiagonal Laplacian matrix with periodic boundary conditions. Time integration was carried out using the Crank-Nicolson method for diffusion terms and explicit Euler method for the reaction terms, implemented via opertor splitting. Linear systems arising from the semi-implicit update steps were successfully solved using a sparse matrix routines from `scipy.sparse` and `scipy.sparse.linalg`.

The five stimulus protocols were: no stimulus (50 cells), front or side stimuli (45 and 15 cells, respectively), global (39 cells) and local-to-global switch (25 cells). For each condition, the fluorescence intensity around the cell edge was recorded over time. The spatial domain was represented from 0 to $2\pi$ with 100 evenly spaced grid points.

Because computing model predictions across all spatial and temporal points for each cell is computationally expensive, we used the average response for each protocol for fitting purposes. The model output was compared to this mean response using a least-squares objective function, and the Differential Evolution algorithm (as in the temporal fitting) was used to optimize the relative rates of diffusion.

To avoid nonphysical or biologically implausible parameter sets, we incorporated penalties into the cost function. For instance, solutions with slower diffusion for inactive Rac relative to active Rac were heavily penalized, based on the known binding of active Rac to the cell membrane. Other penalties were also explored, including a penalty for failure to reverse orientation in response to the local-to-global protocol.

#### 2.4 Stimulus normalization

Later in our model development, we modified the input stimulus $S(x,t)$ to include a normalization factor, dividing it by its integral over the cell edge.

$$S_{\text{new}}(x,t) = \frac{S(x,t)}{\int_\Omega S(x,t)} \tag{6}$$

This normalization reflects the limited range of signalling capacity that biological cells can sense. Mathematically, this normalization helps to balance the model's response to local versus global inputs. Without this adjustment, parameter sets would often fit one protocol well but fail another. Local-fitted models caused overstimulation under global input, leading to a spatial homogeneous solution, while global-fitted ones corresponded to very slow response to local cues. With the normalized stimulus, we achieved consistent fits and lower error across all protocols.

#### 2.5 Results of Spatial Fitting

While Differential Evolution is a global optimization method, the complexity of the parameter space and model outputs in our system makes it unlikely that any single run finds a true global minimum. Moreover, parameter sets with the lowest squared error did not always yield the correct qualitative behaviours, such as proper reversal or sustained polarity. Slight changes in initialization often led to different final fits. For each fitting run, we logged the tested parameter sets, their associated errors, and key model behaviours (e.g., reversal success), allowing us to identify sets that balanced quantitative accuracy with expected biological responses. The parameter set we report here was selected for having near-minimal error while still preserving these key behaviours.

Fitting the WP model was particularly challenging. Even during the temporal fitting (Table 1), several parameters sets failed to converge to biologically meaningful solutions, reflected in the lower success count (58 for WP vs. 87 for WPI). In the spatial setting, this issue persisted. Many WP fits converged to parameter sets that minimized error but led to implausible results, often producing nearly equal diffusion rates for active and inactive Rac. Penalties for missing reversal behaviour were not applied to WP, since it cannot reverse.

The final reported diffusion rates are in units of $\text{rad}^2/\text{s}$, representing effective diffusion along the 1D edge of the 2D projected cell shape. To approximate physical diffusion coefficients in $\mu\text{m}^2/\text{s}$, we used the mean perimeter to motile cells from the reversal experiment dataset (76 µm) to scale the values accordingly. Specifically, since the 1D domain has length $2\pi$, the conversion factor is $(76/2\pi) \approx 146$, which we applied to all fitted values. A summary table of the raw fitted values, their converted counterparts in $\mu\text{m}^2/\text{s}$, and the resulting diffusion rate ratios is given in Table S2.

| Diffusion Coefficient | Ratio (norm. to $D_u$) | rad$^2$/s | $\mu$m$^2$/s |
|---|---|---|---|
| $D_u$ (Active Rac) | 1.00 | 0.080 | 11.705 |
| $D_v$ (Inactive Rac) | 6.40 | 0.510 | 74.617 |
| $D_h$ (Inhibitor) | 0.47 | 0.037 | 5.413 |

Table S2: Fitted diffusion coefficients for Rac model components. Values in µm$^2$/s are computed by rescaling from a $[0, 2\pi]$ domain to a perimeter of 76 µm. Note that this significantly overestimates the true diffusion, as explained below.

However, it is important to keep in mind that these values are "effective rates" of model diffusion along a 1D cell perimeter, based on a 2D fluorescence projection of a 3D phenomenon. The mapping to actual rates of diffusion depend on the geometry. To better understand how these fitted rates compare to real cellular diffusion, we next consider how geometry and dimensionality affect diffusion scaling.

##### 2.5.1 Scaling diffusion

In our simulations, the signalling components are restricted to the cell perimeter, whereas in the real cells, they diffuse on the membrane surface (active form) and in the cell bulk. Here we show how the "effective" rates of diffusion that we found in fitting our model to data (Table XX) correspond to "real rates of diffusion" that would be measured in cells.

In a CPM simulation, the user specifies a target cell area $A_0$ (in pixels), and a target perimeter, $P_0$. As a first approximation, the mean diameter of a simulated cell is

$$d_0 \approx 2\sqrt{A_0/\pi}.$$

The perimeter can be expressed as

$$P_0 = \phi(\pi d_0) = \phi \cdot 2\sqrt{\pi A_0}$$

where $\phi > 0$ is the previously defined "asphericity".

The ratio of the length of the cell perimeter (the domain of the model PDEs) to the cell diameter (1D slice of the domain where actual diffusion takes place) is

$$\frac{P_0}{d_0} = \phi\pi.$$

For our simulations, the asphericity was set to $\phi = 1.4$ to obtain cells resembling neutrophils, so we have a domain ratio that corresponds to

$$\frac{L_1}{L_2} = \frac{P_0}{d_0} \approx 3.14 \cdot 1.4 \approx 4.4$$

It is well known that effective rates of diffusion in domains of size $L_1, L_2$ have the scaling property

$$\frac{D_1}{D_2} \approx \left(\frac{L_1}{L_2}\right)^2 \approx 20.$$

More detailed calculation that represents the 2D geometry of diffusion on the cell surface introduces Bessel functions. This mapping from 1D to 2D also affects the comparison (reducing the “real diffusion” by comparison to the model’s “effective diffusion value” along the perimeter).

Hence, the values fitted for the model imply that actual rates of diffusion are roughly 10-20 fold lower, well in line with values of diffusion for GTPases cited in the literature. For example in [13], typical values are given as 0.1 $\mu m^2$ for diffusion of active GTPase (in the membrane) and 10 $\mu m^2$ for diffusion of the the inactive GTPase (in the bulk).

### 3 Additional Results

#### 3.1 WPI-PIP3 Reproduce Simple Turning Behaviour

We verified that WPI-PIP3 retains the ability to reproduce the basic cell turning behaviours that were originally captured by the WP model. These include the U-turns observed under read stimulation and directional turning in response to one or two stimulation sites on the L and R cell sides. These simple responses are previously shown for the WP model in Figure V of the main text.

Here, we repeat those protocols using the WPI-PIP3 models with its best-fit parameters. As shown in Figure S3, the model produces qualitatively correct turning behaviour: In the rear-stimulation, most of the time the cell makes a U-turn and then moves in the opposite direction. In the one-sided stimulus, the model cell turns towards the stimulus direction, while in the two-sided stimuli, the cell randomly selects a “winning” direction, as seen in the experiments.

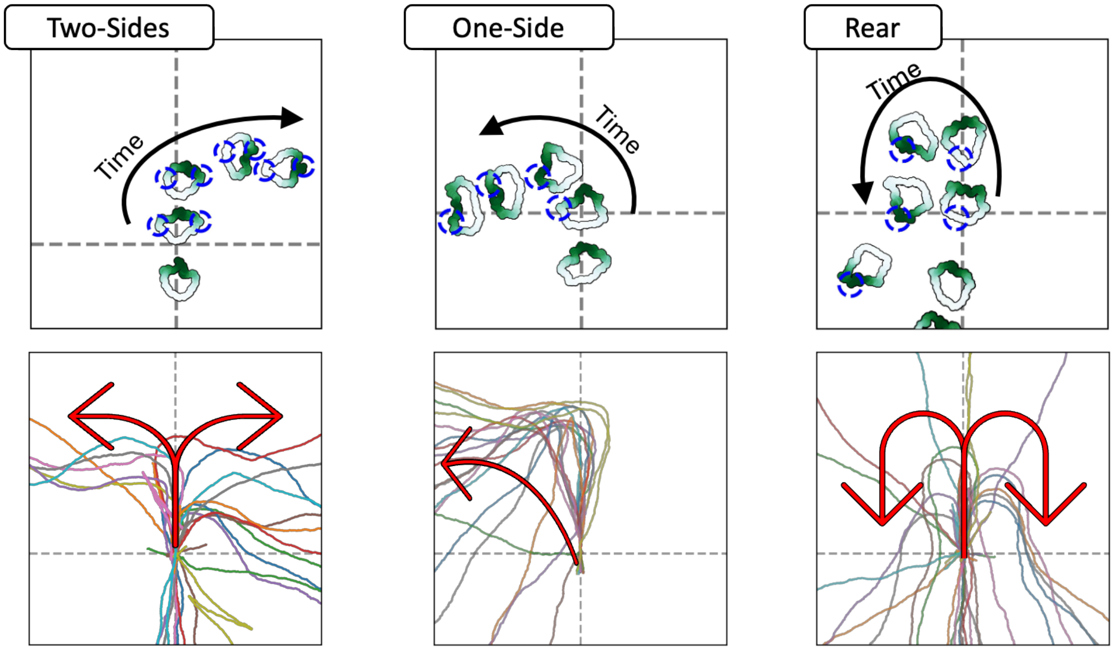


Figure S3: Each column shows results from a distinct stimulus: two-sided stimulation (left), one-sided (middle), and rear stimulation (right). Top row: Representative snapshots from simulated cell trajectories over time. Active Rac is shown in green, light stimulus in blue. Bottom row: Superimposed tracks of 20 simulated cells in response to each stimulus. Red arrows indicate typical direction of turning.The WPI-PIP3 model captures the expected behaviour in each case: random left/right choice under two-sided stimulation, directional turning under one-sided stimulation, and a U-turn under rear stimulation.

#### 3.2 Model Responses to Noisy Chemical Gradients

To assess the robustness of the distinct models (WP, WPI, WPI-PIP3) under more biologically realistic conditions, we simulated cells in a chemical gradient similar to Figure XV. This time, we introduced Gaussian noise with increasing variances to the chemical concentration and did not reverse the gradient at any point.

For each model, we ran 20 simulations (with random initial orientations) in a noise-free gradient and two levels of noise (variance of 0.4 and 0.6, here referred to as medium and high noise intensity, respectively), and recorded the average direction of the Rac peak from $t = 100$ to $t = 200$. The results are summarized as rose-plots in Figure S4, where each subplot shows the distribution of averaged polarization directions for increasing noise intensity (left to right) and across the three models (top to bottom).

As expected, in the absence of noise, all models aligned effectively with the gradient. However, as noise increases, WP quickly loses directional accuracy. WPI performed better, but its alignment was still impaired at higher noise levels. The WPI-PIP3 model consistently outperformed the others, preserving polarity alignment even under high noise. This supports the hypothesis that

PIP3 allows the cell to smooth out noisy input signals before they affect Rac activation.

To validate that the success of the WPI-PIP3 model requires the combined action of both the inhibitor and PIP3, we tested a fourth model variant containing only the WP mechanism and PIP3, but no inhibitor (WP-PIP3).

Figure S5 shows the predicted response of this model variant to the same noisy gradients previously tested. As noise increases, the model rapidly loses its ability to align with the gradient. Despite the smoothing effect of PIP3, the absence of a local inhibitor impairs the cell's ability to detect a weak noisy gradient. This result supports the conclusion that both components are required for robust chemotaxis in complex environments.

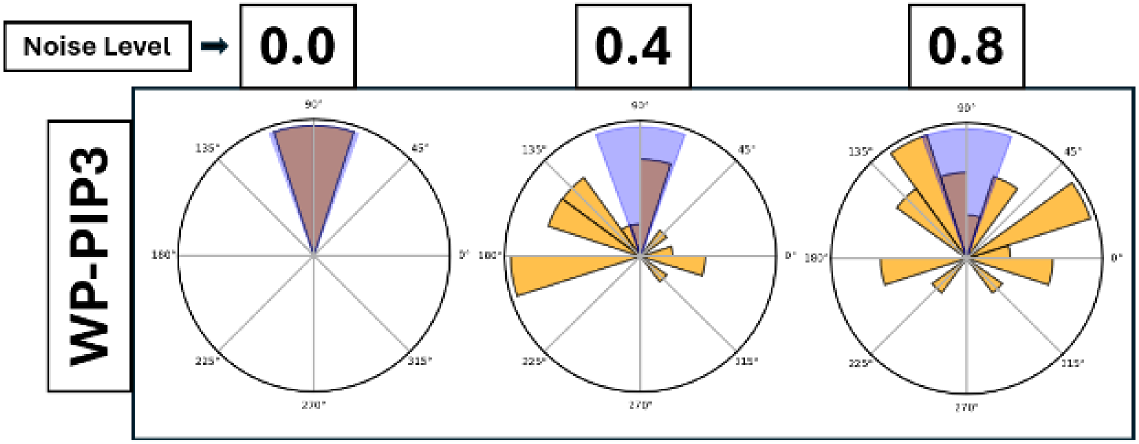


Figure S5: As in Figure S4, but for the model with no inhibitor (WP-PIP3). Each rose plot shows the distribution of averaged Rac polarity directions in cells stimulated with a fixed northward gradient at various levels of noise (variance 0.0, 0.4, 0.8). The models align well in the absence of noise, but its performance degrades rapidly under increasing noise, demonstrating the role of the Rac-inhibitor in robust gradient sensing.

### References

[1] Jason P Town and Orion D Weiner. Local negative feedback of rac activity at the leading edge underlies a pilot pseudopod-like program for amoeboid cell guidance. *PLoS Biology*, 21(9):e3002307, 2023.

[2] Yoichiro Mori, Alexandra Jilkine, and Leah Edelstein-Keshet. Wave-pinning and cell polarity from a bistable reaction-diffusion system. *Biophysical journal*, 94(9):3684–3697, 2008.

[3] Hans Meinhardt. Orientation of chemotactic cells and growth cones: models and mechanisms. *Journal of cell science*, 112(17):2867–2874, 1999.

[4] M. P. Neilson, J. A. Mackenzie, S. D. Webb, and R. H. Insall. Modeling cell movement and chemotaxis using pseudopod-based feedback. *SIAM Journal on Scientific Computing*, 33(3):1035–1057, 2011.

H

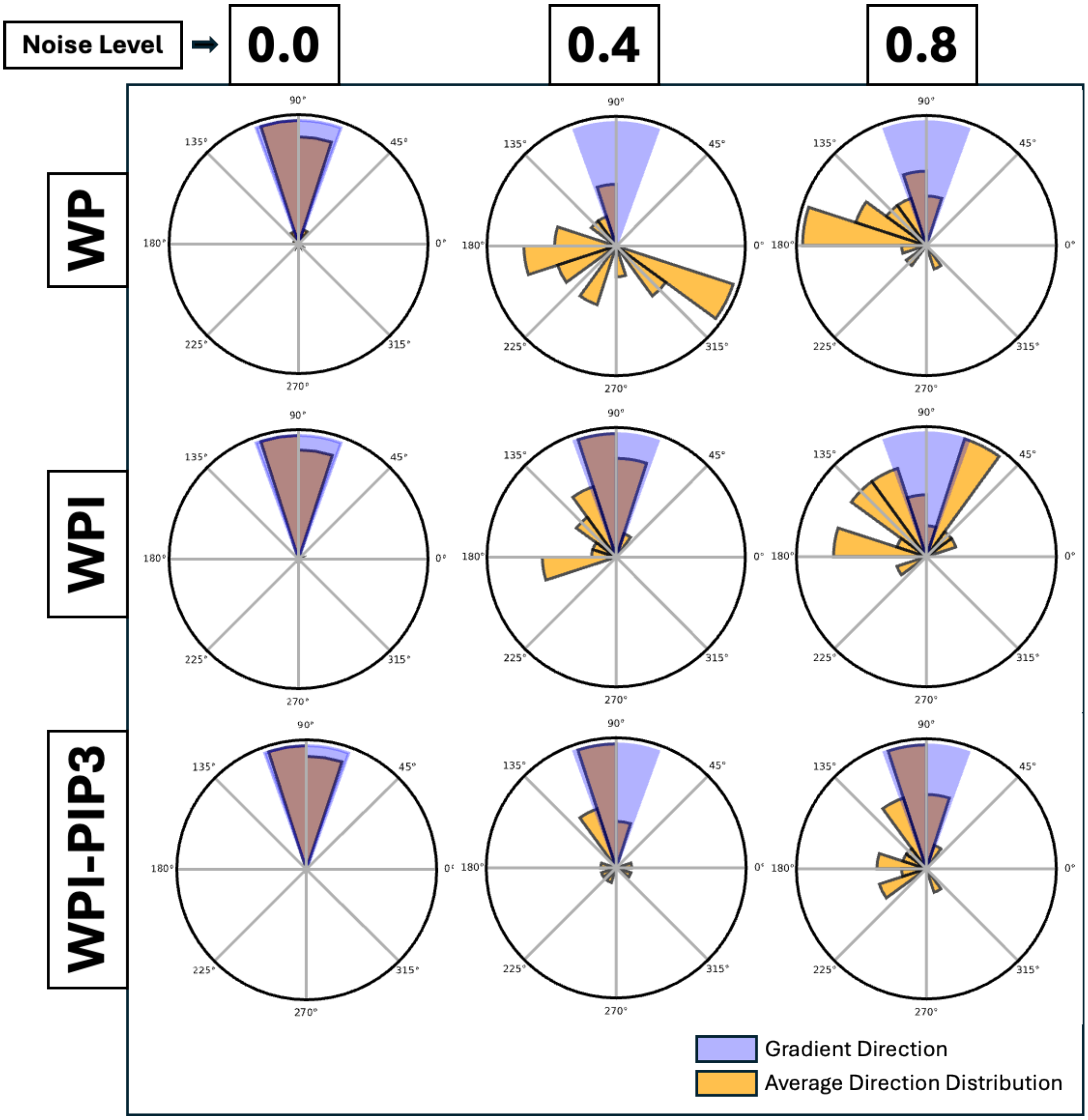


Figure S4: Model responses to noisy chemical gradients. Each row corresponds to a model (top to bottom: WP, WPI, WPI-PIP3), and each column represents a level of noise added to the gradient (left: none, middle: moderate, right: high). All models align well with a gradient without noise, but only the WPI-PIP3 allows the cell to sense highly noisy gradients.

[5] Jörn Starruß, Walter De Back, Lutz Brusch, and Andreas Deutsch. Morpheus: a user-friendly modeling environment for multiscale and multicellular systems biology. *Bioinformatics*, 30(9):1331–1332, 2014.

[6] François Graner and James A Glazier. Simulation of biological cell sorting using a two-dimensional extended potts model. *Physical review letters*, 69(13):2013, 1992.

[7] Athanasius FM Marée, Alexandra Jilkine, Adriana Dawes, Verônica A Grieneisen, and Leah Edelstein-Keshet. Polarization and movement of keratocytes: a multiscale modelling approach. *Bulletin of mathematical biology*, 68:1169–1211, 2006.

[8] Elisabeth G Rens and Leah Edelstein-Keshet. Cellular tango: how extracellular matrix adhesion choreographs rac-rho signaling and cell movement. *Physical biology*, 18(6):066005, 2021.

[9] Ioana Niculescu, Johannes Textor, and Rob J De Boer. Crawling and gliding: a computational model for shape-driven cell migration. *PLoS computational biology*, 11(10):e1004280, 2015.

[10] Alexandra Jilkine and Leah Edelstein-Keshet. A comparison of mathematical models for polarization of single eukaryotic cells in response to guided cues. *PLoS computational biology*, 7(4):e1001121, 2011.

[11] Hao Chang and Andre Levchenko. Adaptive molecular networks controlling chemotactic migration: dynamic inputs and selection of the network architecture. *Philosophical Transactions of the Royal Society B: Biological Sciences*, 368(1629):20130117, 2013.

[12] Kenneth Price, Rainer M Storn, and Jouni A Lampinen. *Differential evolution: a practical approach to global optimization*. Springer Science & Business Media, 2006.

[13] Marten Postma, Leonard Bosgraaf, Harriët M Loovers, and Peter JM Van Haastert. Chemotaxis: signalling modules join hands at front and tail. *EMBO reports*, 5(1):35–40, 2004.
